# A Microphysiological HHT-on-a-Chip Platform Recapitulates Patient Vascular Lesions

**DOI:** 10.1101/2024.03.11.584490

**Authors:** Jennifer S. Fang, Christopher J. Hatch, Jillian Andrejecsk, William Van Trigt, Damie J. Juat, Yu-Hsi Chen, Satomi Matsumoto, Abe P. Lee, Christopher C.W. Hughes

## Abstract

Hereditary Hemorrhagic Telangiectasia (HHT) is a rare congenital disease in which fragile vascular malformations focally develop in multiple organs. These can be small (telangiectasias) or large (arteriovenous malformations, AVMs) and may rupture leading to frequent, uncontrolled bleeding. There are few treatment options and no cure for HHT. Most HHT patients are heterozygous for loss-of-function mutations for Endoglin (ENG) or Alk1 (ACVRL1), however, why loss of these genes manifests as vascular malformations remains poorly understood. To complement ongoing work in animal models, we have developed a microphysiological system model of HHT. Based on our existing vessel-on-a-chip (VMO) platform, our fully human cell-based HHT-VMO recapitulates HHT patient vascular lesions. Using inducible *ACVRL1* (Alk1)-knockdown, we control timing and extent of endogenous Alk1 expression in primary human endothelial cells (EC) in the HHT-VMO. HHT-VMO vascular lesions develop over several days, and are dependent upon timing of Alk1 knockdown. Interestingly, in chimera experiments AVM-like lesions can be comprised of both Alk1-intact and Alk1-deficient EC, suggesting possible cell non-autonomous effects. Single cell RNA sequencing data are consistent with microvessel pruning/regression as contributing to AVM formation, while loss of PDGFB expression implicates mural cell recruitment. Finally, lesion formation is blocked by the VEGFR inhibitor pazopanib, mirroring the positive effects of this drug in patients. In summary, we have developed a novel HHT-on-a-chip model that faithfully reproduces HHT patient lesions and that is sensitive to a treatment effective in patients. The VMO-HHT can be used to better understand HHT disease biology and identify potential new HHT drugs.

**Significance:** This manuscript describes development of an organ-on-a-chip model of Hereditary Hemorrhagic Telangiectasia (HHT), a rare genetic disease involving development of vascular malformations. Our VMO-HHT model produces vascular malformations similar to those seen in human HHT patients, including small (telangiectasias) and large (arteriovenous malformations) lesions. We show that VMO-HHT lesions are sensitive to a drug, pazopanib, that appears to be effective in HHT human patients. We further use the VMO-HHT platform to demonstrate that there is a critical window during vessel formation in which the HHT gene, Alk1, is required to prevent vascular malformation. Lastly, we show that lesions in the VMO-HHT model are comprised of both Alk1-deficient and Alk1-intact endothelial cells.

## Introduction

Hereditary Hemorrhagic Telangiectasia (HHT) is a rare congenital disease that impacts 1 in 5000 people and is characterized by the sporadic development of vascular malformations.^1,2^ These disorganized lesions include small, dilated, and fragile tangles of vessels called telangiectasias^3^ that affect skin and mucosa, especially in the nose, where their rupture leads to frequent and uncontrolled epistaxis (nosebleeds) that greatly impacts patient quality-of-life.^4,5^ Many HHT patients also develop larger arteriovenous malformations (AVM) – enlarged arteriolar-to-venous shunts^6^ – within their gut, lung, liver, and brain that can compromise tissue perfusion leading to organ damage and eventual failure.^4^ HHT is sub-divided into disease subtypes based on patient gene mutation status. Greater than 90% of HHT patients inherit loss-of-function mutations affecting either *ENG* (HHT1), which encodes Endoglin, or *ACVRL1* (HHT2), which encodes Activin-like Kinase receptor 1 (Alk1).^4^ Eng and Alk1 are endothelium-expressed cell surface TGF-β superfamily co-receptors, with Alk1 being the signal transducing component, and mutation of either affects Alk1-driven signaling through downstream Smads. Specifically, Alk1 receptor activation phospho-activates the Smad1/5 complex, which then binds to Smad4. The activated Smad complex translocates into the nucleus to regulate target gene expression.^7^ A rare variant of HHT – juvenile polyposis-HHT (JP-HHT)^8^ – accounts for ∼2% of HHT cases and has been mapped to the *SMAD4* gene, a key integrator in the Smad pathway.^8–10^ Importantly, patients are heterozygous for Alk1, Eng, or Smad4 mutations, with lesions being thought to occur as a result of somatic mutation of the residual allele.^11–13^ A recent study that used deep sequencing of isolated HHT patient lesions has confirmed this hypothesis, but with the surprising additional observation that many of the lesions in the same patient are of mixed composition, including both heterozygous and subpopulations of homozygously-mutated endothelial cells (EC) containing different second hits – suggesting multiple loss-of-heterozygosity (LoH) events.^12^ Relatedly, mouse studies have also shown that homozygous deletion of Endoglin in only a small subpopulation of EC is sufficient to induce AVMs when combined with a focal pro-angiogenic stimulus.^14^

The relevant ligands for the Alk1/Eng receptor complex are BMP9 and BMP10,^15^ although other TGF-β ligands are also capable of binding at much higher concentrations.^15^ Of these, BMP9 is present in blood at concentrations well above its binding affinity for the Alk1 receptor, suggesting that there is tonic signaling through this pathway in established vasculature, possibly to maintain EC quiescence.^16,17^

Unfortunately, there is currently no cure for HHT and few treatment options, due in part to the challenges of modeling HHT vascular malformations in the preclinical setting. Most research on the pathogenesis of vascular malformations has been conducted in genetically-modified mouse models wherein Alk1, Eng, or Smad4 are homozygously ablated, either globally or only in vascular endothelium.^18^ Although these models have provided important insights into HHT disease biology,^9,10,14,19–21^ AVMs occur only sporadically and can be difficult to identify and monitor in real-time. Several key insights have also been provided by zebrafish models, including the insight that Alk1 is a flow-sensitive gene and that dysregulated flow-directed EC migration may contribute to vascular malformation in HHT.^22–24^ In this study, we use a microphysiological system (MPS) platform to generate a human cell-based *in vitro* microphysiological model of vascular malformations in HHT that can complement and enhance research in animal models.

MPS -- or Organ-on-a-chip (OoaC) -- refers to the engineering of artificial microtissues in a three-dimensional *in vitro* microenvironment that captures the microphysiology of native, intact tissue.^25,26^ MPS models enable reproducible and high-throughput engineering of complex and physiological microtissues in an *in vitro* setting. They are thus powerful new tools for understanding healthy and diseased tissues that will help in translational research by supporting both compound library drug screens and personalized medicine applications.^25,27^

We previously developed a human vessel-on-a-chip (Vascularized Micro-Organ, VMO) platform that combines primary human endothelial and perivascular cell types cultured in a three-dimensional hydrogel and in the presence of interstitial flow.^27–30^ Under these conditions, EC self-organize over the course of 4-5 days into a lumenized microvascular network that is perfused by a gravity-driven blood substitute. We have previously used the VMO platform to study vascularized micro-tumors,^28,31–33^ as well as to model healthy tissue-vascular interactions.^29,34^ In this study, we describe an adapted VMO platform that allows for the development of disorganized vessel lesions with the architectural hallmarks of HHT telangiectasias and AVMs. We call this novel HHT-on-a-chip microphysiological platform the HHT-VMO and show that it can be used to provide valuable insights into HHT disease biology and to evaluate drugs for their therapeutic potential in HHT.

## Results

### Alk1-deficient EC form hyperdense microvasculature in vitro

As noted above, HHT patients are germline heterozygous for loss-of-function mutations that affect either Alk1 (*ACVRL1*) or its co-receptor Eng (*ENG*).^4^ To generate EC lacking intact Alk1 signaling, mimicking the LoH events thought to occur in lesions, we initially pursued a transient silencing RNA (siRNA) approach to target endogenous *ACVRL1* (si-*ACVRL1*) transcript. This achieved a near-complete knockdown of Alk1 mRNA that persisted for at least 7 days in primary human umbilical vein EC (HUVEC), but this knockdown fully recovered to control levels by day 12 (**Supplemental Figure 1A**). We seeded EC transfected with si-*ACVRL1* (or scrambled si-Ctrl) into our previously described VMO platform^28,29,31,35^ at equal starting cell densities (**Supplemental Figure 1B-C**) and observed that microvasculature that formed from si-*ACVRL1* EC were hyperdense relative to si-Ctrl (**Supplemental Figure 1D-I**). We observed a similar phenotype with siRNA knockdown of *ENG* (**Supplemental Figure 2**), suggesting that this hyperdense phenotype arises from loss of signaling through the Alk1/Eng HHT-associated signaling pathway.

Next, to generate stable Alk1 knockdown that persists for longer than 7 days, we used lentiviral transduction to introduce an Isopropyl ß-D-1-thiogalactopyranoside (IPTG)-sensitive *ACVRL1*-targeting short hairpin RNA (shRNA) into primary HUVEC at >85% transduction efficiency (**Supplemental Figure 3A-B**). IPTG is an allolactose mimetic frequently used as an inducing agent to activate tunable transcription of genes under control of the *lac* operon, such as in our IPTG-inducible sh-*ACVRL1* system. In resulting sh-*ACVRL1* EC, addition of IPTG into the perfusion medium induced dose-dependent Alk1-deficiency (**Supplemental Figure 3C**). 3mM IPTG produced >90% *ACVRL1* mRNA knockdown, and transduction of the construct into EC was stable as evidenced by a persistently high-efficiency of *ACVRL1* gene knockdown at day 14 in response to IPTG (**Figure 1A**). Importantly, we tested two independent shRNA sequences (Table 1) and results were indistinguishable (data not shown). Consequently, we used them interchangeably in the reported studies. Alk1 protein expression was also reduced in sh-*ACVRL1* EC in response to IPTG (**Figure 1B**). Consistent with our initial experiments using siRNA (i.e. short-term Alk1 or Eng knockdown, **Supplemental Figures 1-2**), IPTG-treated (i.e. Alk1-deficient) sh-*ACVRL1* EC formed hyperdense microvasculature in our published VMO^28,32^ platform (schematic in **Supplemental Figure 4**) compared to untreated controls (**Figure 1C**). This observed increase in microvessel density was quantified in FIJI/ImageJ as a significantly increased vessel area (**Figure 1D**), increased total vessel length (**Figure 1E**), increased branchpoint number (**Figure 1F**), and a decreased mean lacunarity (**Figure 1G**). At later times overall vessel length was similar in control and knockdown, likely due to the physical constraints of the system. Perhaps related to this, vessel diameter was increased at later times **(Figure 1H)**. Importantly, IPTG alone had no effect on microvascular network formation in the VMO (**Supplemental Figure 5A-D**). Interestingly, even though knockdown of Alk1 produced higher microvessel density and more enlarged vessels in the VMO, vessel network architecture remained relatively ordered in appearance, with no obvious development of disorganized focal lesions that resembled AVMs.

**Figure 1.**
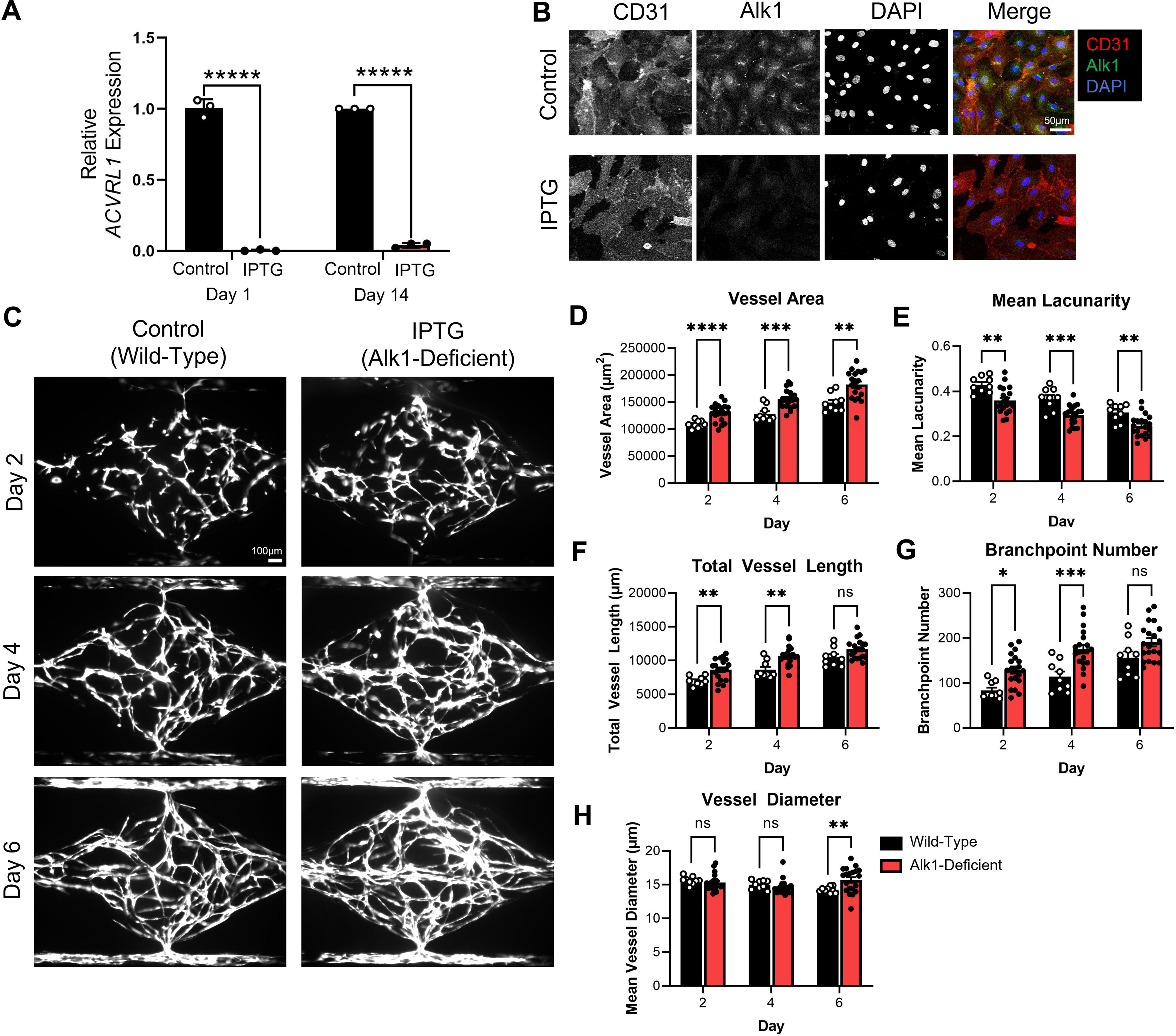
Alk1-deficient EC (via IPTG-sensitive shRNA) form hyperdense microvasculature in VMO. **A)** IPTG eliminates >95% endogenous Alk1 mRNA expression in human EC at day 1 and day 14 post-transduction (n=3 for each group). **B)** Endogenous Alk1 protein expression was also reduced in EC. **C)** IPTG-treated Alk1-deficient EC form hyperdense microvasculature compared to untreated (i.e., no IPTG) Alk1-intact control EC. This was associated with increased vessel area **(D)**, increased total vessel length **(E)**, increased branchpoint number **(F)**, and decreased mean lacunarity **(G).** Mean vessel diameter was increased but only at the later timepoint **(H)**. (n=9 Control, n=20 IPTG). (Statistics: * - p < 0.05, ** - p < 0.01, *** - p < 0.001, **** - p < 0.0001, ***** - p < 0.00001)

**Figure 2.**
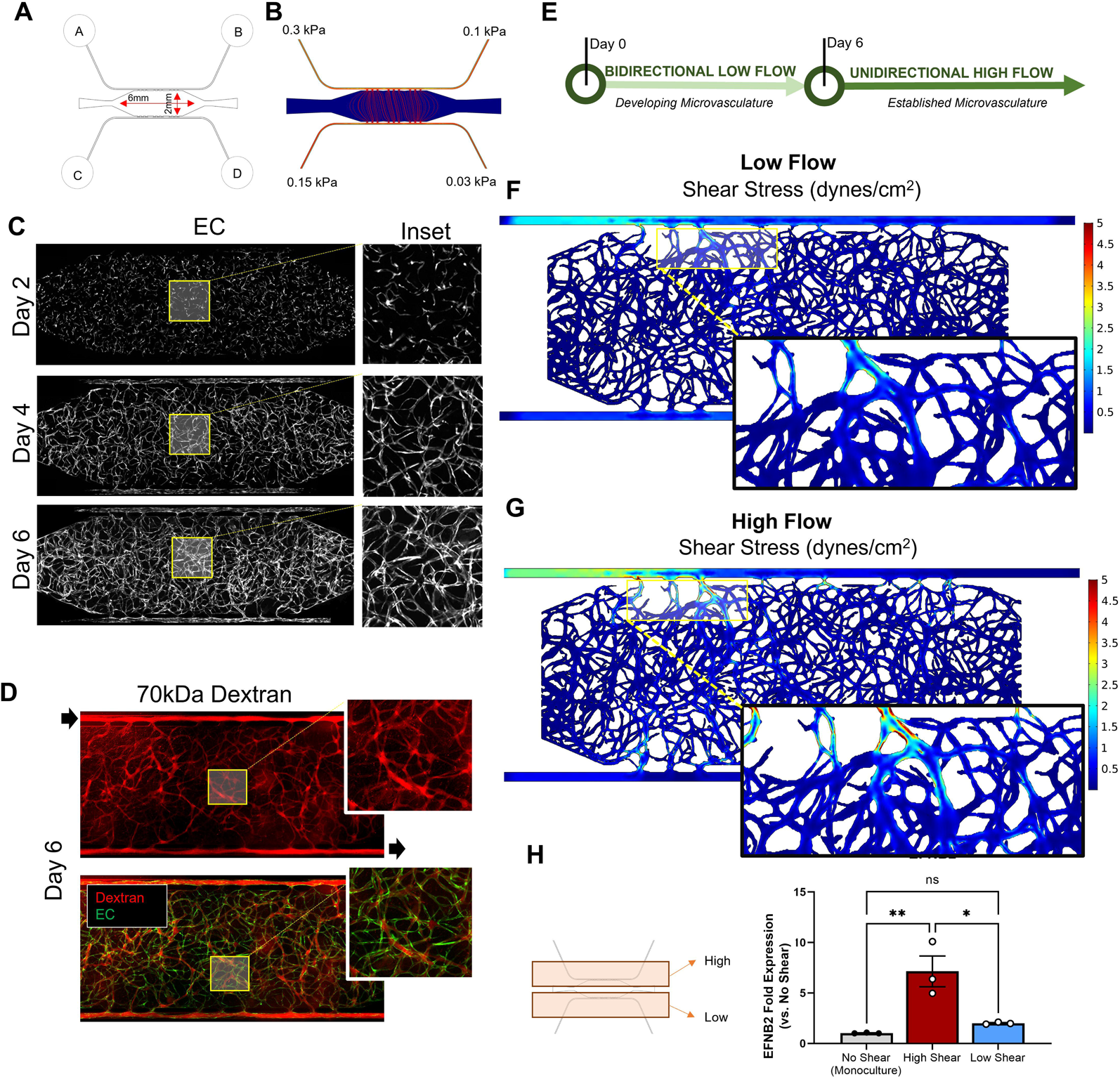
Modified microfluidic vessel-on-a-chip platform (HHT-VMO) supports formation of a perfused microvascular network under physiological shear. **A)** Redesign of VMO platform to enlarge central tissue chambers and media reservoirs and to uncouple microfluidic channels. Flow is from A to B alone the “artery” and C to D along the “vein”. Higher pressure on the arterial side drives net flow from A to D through the vasculature. **B)** COMSOL modeling shows pressure differentials and interstitial flow lines prior to vessel establishment. **C)** Seeding of fluorescent reporter-expressing primary human EC in the HHT-VMO under interstitial flow leads to the formation of perfused microvascular network over ∼6 days. **D)** Perfused microvessels are visualized by addition of a 70kDa fluorescent dextran (red) to the media reservoirs. EC are green. **E)** Two-step flow protocol for inducing an established microvasculature. F, G) *In silico* modeling of low and high flow settings produces regions of more physiological fluid shear stress (∼5 dynes/cm^2^), particularly around the communication pores in the upper (arterial) media channel (insets). **H)** Following 2 days exposure to high flow, EC in the high and low shear regions were isolated for qPCR analysis. EC in the high shear zone acquire expression of the arterial marker ephrin B2 (*EFNB2*). (Statistics: ** - p < 0.01)

**Table 1.**
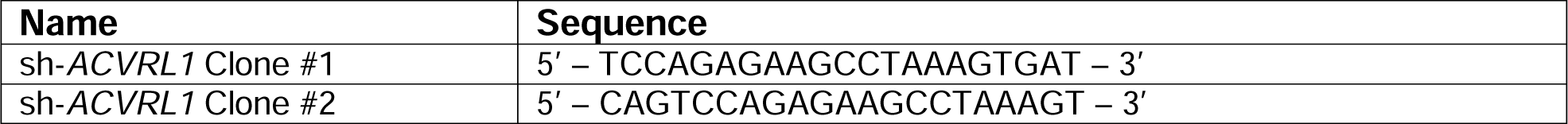
sh-*ACVRL1* Clones.

### HHT-VMO platform supports physiological flow at arteriolar levels

In mouse models of HHT, vascular malformations appear in the neonatal retina in regions associated with higher fluid shear stress, whereas at lower levels of shear, vascular overgrowth (telangiectasia-like) is seen,^36^ suggesting that hemodynamic flow is a critical initiating signal for the development of these lesions. Using *in silico* approaches, we modeled flow in our VMO platform and found that a maximal fluid velocity of 2μm/s, producing a maximal shear stress of <0.5 dynes/cm^2^ (**Supplemental Figure 6**). These values are significantly lower than the flow velocity and shear stress values calculated for the regions predisposed to shunt formation in the retinal vasculature of neonatal mice.^37^

To generate a platform with a more physiologic range of intravascular flow and fluid shear stress, we adapted our base VMO (**Supplemental Figure 4**) by uncoupling the upper and lower microfluidic channels and enlarging the central tissue chamber and media reservoirs (**Figure 2A**). This indeed led to higher levels of interstitial flow (**Figure 2B**), but still in the range necessary to induce formation of a perfused microvascular network. Network formation took 6-8 days following initial cell seeding (**Figure 2C**), consistent with how the microvasculature develops in our published VMO models.^28,29,31,35^ Once vessel formation was complete, addition of 70kDa fluorescent dextran to the media reservoirs marked lumenized microvessels and revealed that vessels are perfused and that flow is *only* through vessels (**Figure 2D**). To model the normal developmental process, we next developed a two-step flow protocol to support the development of a perfused microvasculature under physiologic flow, wherein EC are initially exposed to bidirectional low flow for the first 6 days, mirroring the hemodynamic conditions of developing microvessels. After 6 - 8 days, networks were switched to unidirectional high (physiological) flow, consistent with the conditions present upon establishment of a systemic circulation *in vivo* (**Figure 2E**). *In silico* modeling of low-(**Figure 2F**) and high-flow (**Figure 2G**) conditions confirms that they support physiologic fluid shear stress levels of ∼1 dynes/cm^2^ and ∼5 dynes/cm^2^, respectively. Importantly, we observed increased expression of the endothelial-specific arterial marker Ephrin B2 (*EFNB2*) only in regions of high shear stress following 2 days of high flow in the HHT-VMO platform (**Figure 2H**). In parallel, perivascular cells in the high shear stress region of the microvasculature also acquire expression of smooth muscle _α_-Actin (SMA) (**Supplemental Figure 7**), another indicator of vessel arterialization under these conditions.

### Alk1-deficient EC in the modified platform generate disorganized microvascular networks with lesions reminiscent of HHT patient telangiectasias

We generated EC transduced with the IPTG-sensitive sh-*ACVRL1* construct, which were then seeded into the HHT-VMO platform. IPTG was added to the medium beginning at day 0 and maintained throughout the timecourse of the experiment to induce knockdown of endogenous Alk1 expression. Wild-type (control) networks were not exposed to IPTG, thereby leaving Alk1 expression intact. Consistent with our observations in the original (low-flow) VMO platform (**Figure 1**), IPTG-treated (Alk1-deficient) EC in the modified platform also formed hyperdense networks compared to wild-type controls when the shear was at the lower end of physiologic levels (**Figure 3A**). This was quantified as a significantly increased vessel area, total vessel length and branchpoint number, as well as significantly decreased mean lacunarity, by day 6 (**Figure 3B-E**). When networks were switched to high flow in the presence of IPTG, we observed microvascular structures that progressively grew over time (**Figure 3F**), resulting in enlarged and dilated focal lesions that varied in their presentation from dense clusters or tangles to enlarged loop-like structures (**Figure 3G**). This disorganized architecture is reminiscent of HHT patient telangiectasias, which are also highly variable and typically present in the nasal septum as “spots”, “loops”, “spiders”, or “raspberry”-like tangles of disorganized microvasculature.^3^

**Figure 3.**
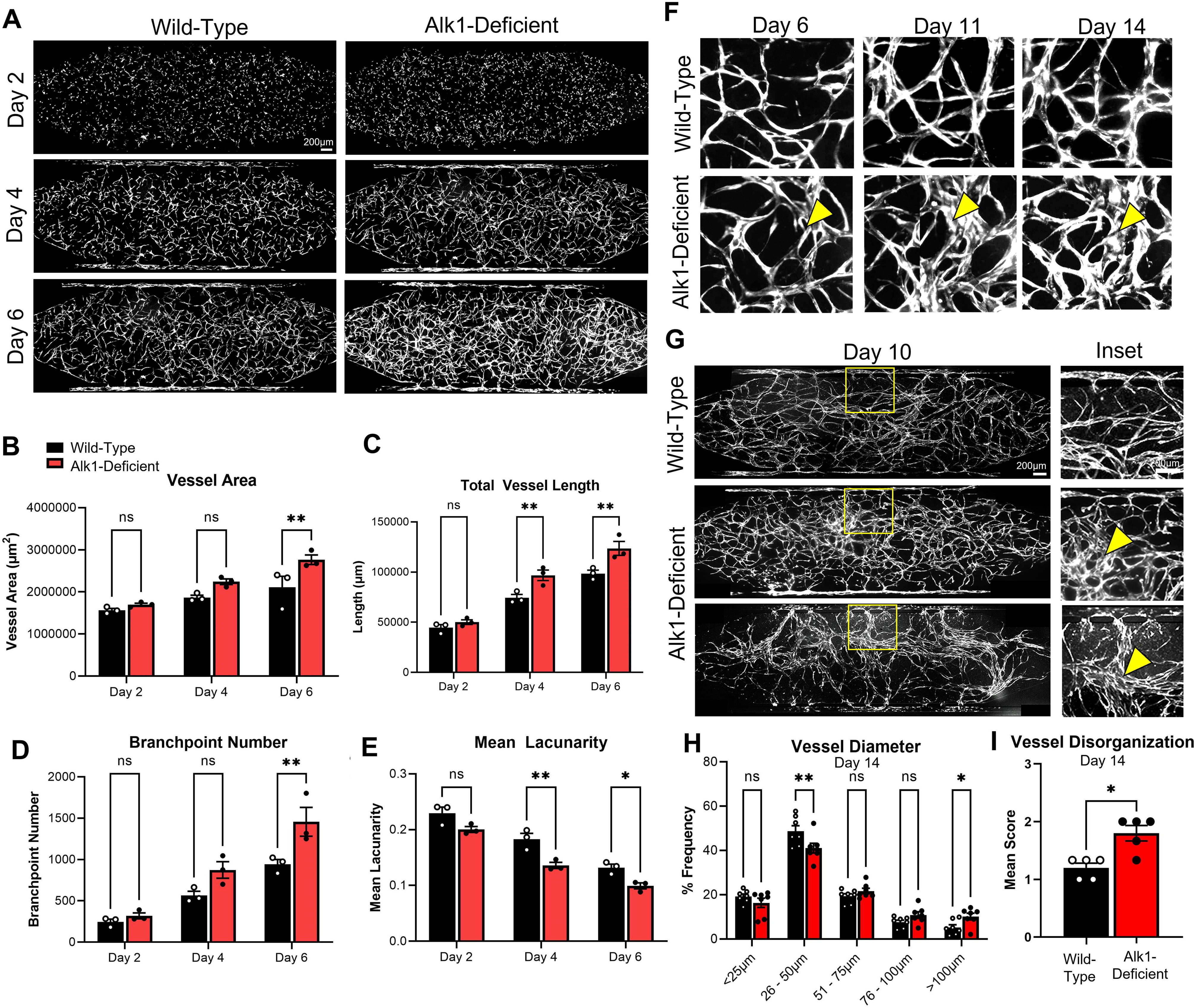
Alk1-deficient EC form hyperdense and disorganized microvasculature in the HHT-VMO platform. **A)** Compared to control networks, IPTG-treated networks appear hyperdense during initial network formation, quantified by vessel morphometry as significant increases in vessel area **(B)**, total vessel length **(C)**, and branchpoint number **(D)**. Mean lacunarity was decreased by day 6 **(E)**. **F)** IPTG-treated networks contained structures that continued to dilate with additional time in culture (yellow arrowhead) **(G)** By day 10 we observed networks with focally dilated lesions (middle), or a complete collapse of the microvascular network to a few dominant vessels (bottom). **(H)** At day 14, IPTG-induced Alk1-knockdown increased the frequency of vessels with a larger diameter. **I)** Semi-quantitative “vascular disorganization score” was increased in the absence of Alk1. (Statistics: * - p < 0.05, ** - p < 0.01).

Consistent with the observed appearance of dilated telangiectasia-like lesions with addition of circulating IPTG, morphometric analysis of the entire microvascular network revealed a significant increase in the frequency of vessels with a diameter >50 µm and a concomitant decrease in the frequency of vessels with a diameter <50 µm (**Figure 3H**). Using an investigator-blinded semi-quantitative scoring approach (see **Materials and Methods**), we also detected a significant increase in vessel disorganization score in IPTG-treated networks (**Figure 3I**). Importantly, there was no effect on vessel network appearance when wild-type EC (**Supplemental Figure 5E-H**) were treated with IPTG in the platform.

### Lesion formation only occurs when Alk1 is lost early in vessel development

To assess when Alk1 expression is most critical during vessel development, we seeded sh-*ACVRL1* EC into the HHT-VMO and applied IPTG to the perfusion medium prior to network formation (day 0, as described above), during active network formation (day 3), or after the vasculature was largely established (day 8). We then assessed network morphology at days 10 and 14, to allow a minimum of 6 days in each group for lesions to develop after onset of IPTG; these timepoints were selected based on our earlier studies (**Figure 3**) showing that lesions became obvious by day 6 when IPTG was administered at day 0. Remarkably, only the administration of IPTG at day 0 or day 3, but not at day 8, led to the appearance of disorganized vascular lesions (**Figure 4**). This indicates that there is a critical early window during vessel network development when the absence of Alk1 can trigger lesion formation. In the HHT-VMO system, this window is prior to 8 days. These findings are consistent with work from the Oh lab^14,19,20^, which has used mouse models of HHT to show that loss of Alk1 expression drives lesion formation in the early phases of an angiogenic response, whereas loss of Alk1 in stable vessels in the absence (or neutralization) of pro-angiogenic signaling does not produce a phenotype.

**Figure 4.**
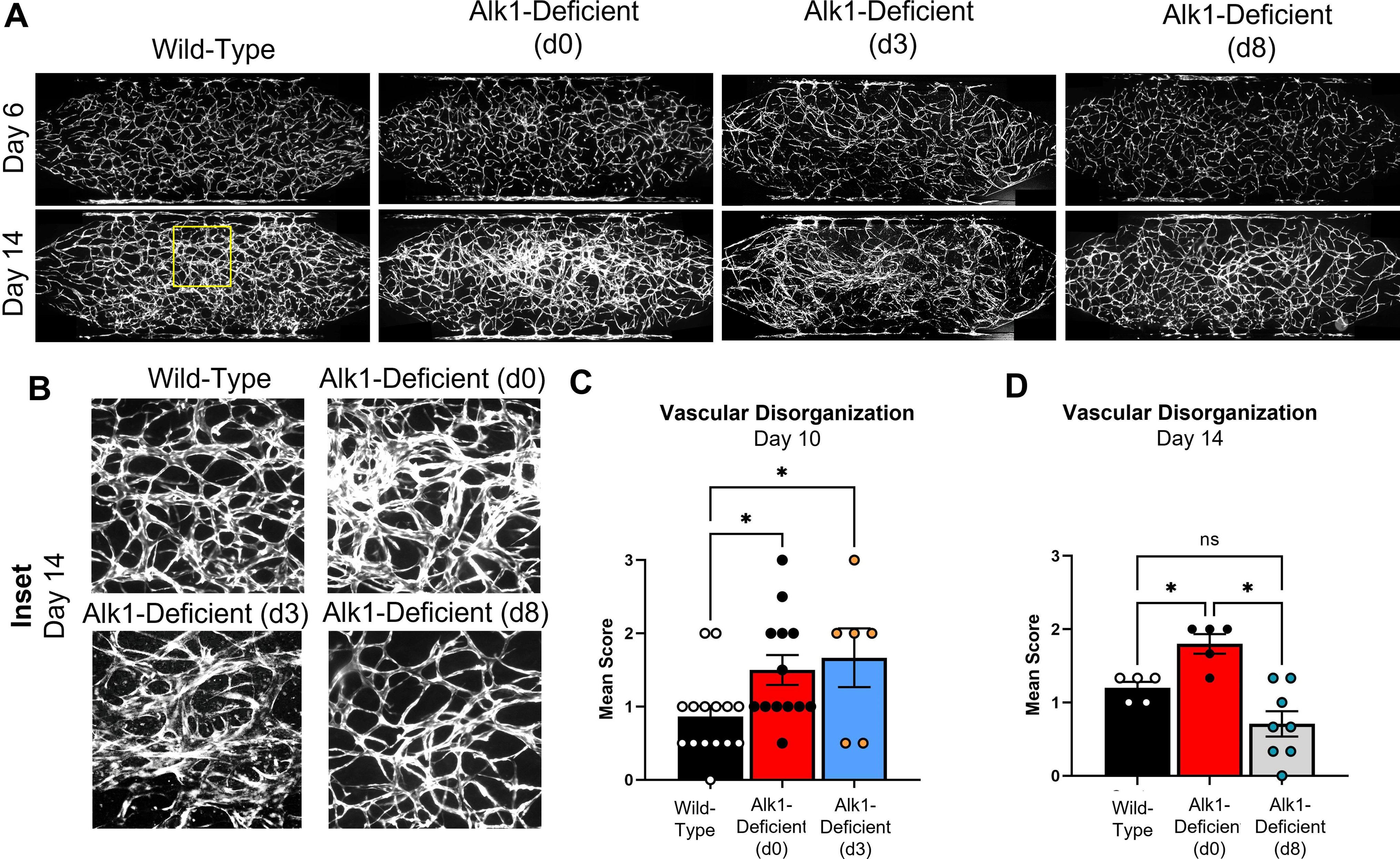
Alk1 knockdown in HHT-VMO during (but not after) network formation results in microvascular lesions. **A)** IPTG was administered to initiate Alk1 knockdown at days 0 or 3, during microvascular network formation, or at day 8 once the microvascular network is established. **B)** Alk1 knockdown at day 0 or day 3, but not day 8, induced the appearance of disorganized microvascular lesions. All insets reflect the same region in the upper middle of their respective networks as denoted by the yellow box in panel A. **C)** Vascular disorganization scores at day 10 show similarly elevated disorganization whether Alk1 is knocked down at day 0 or day 3. **D)** Vascular disorganization scores at day 14 show elevated disorganization scores only in networks treated with IPTG at day 0. (Note, the data in Figure 3I are a subset of these data). (Statistics: * - p < 0.05).

### Pazopanib prevents lesion formation

Pazopanib is a VEGFR inhibitor that has been shown to improve bleeding symptoms in HHT patients,^38,39^ although the drug’s effects on underlying vascular malformations in HHT patients remains uncertain. To assess whether HHT-VMO lesions could be prevented by application of pazopanib, we seeded shACVRL1 EC into the HHT-VMO and added IPTG from day 0. We then added 25nM (*data not shown*) or 100nM pazopanib (or DMSO vehicle) to the perfusing medium beginning at day 3, when our data indicate that Alk1 remains necessary to prevent vascular lesions (**Figure 4**). Consistent with the clinical data showing bleeding and anemia improvement with clinical administration of pazopanib,^38,39^ we observed an inhibition of lesion formation when pazopanib was added to IPTG-treated networks (**Figure 5**) with resultant vascular networks being barely distinguishable from wild-type networks.

**Figure 5.**
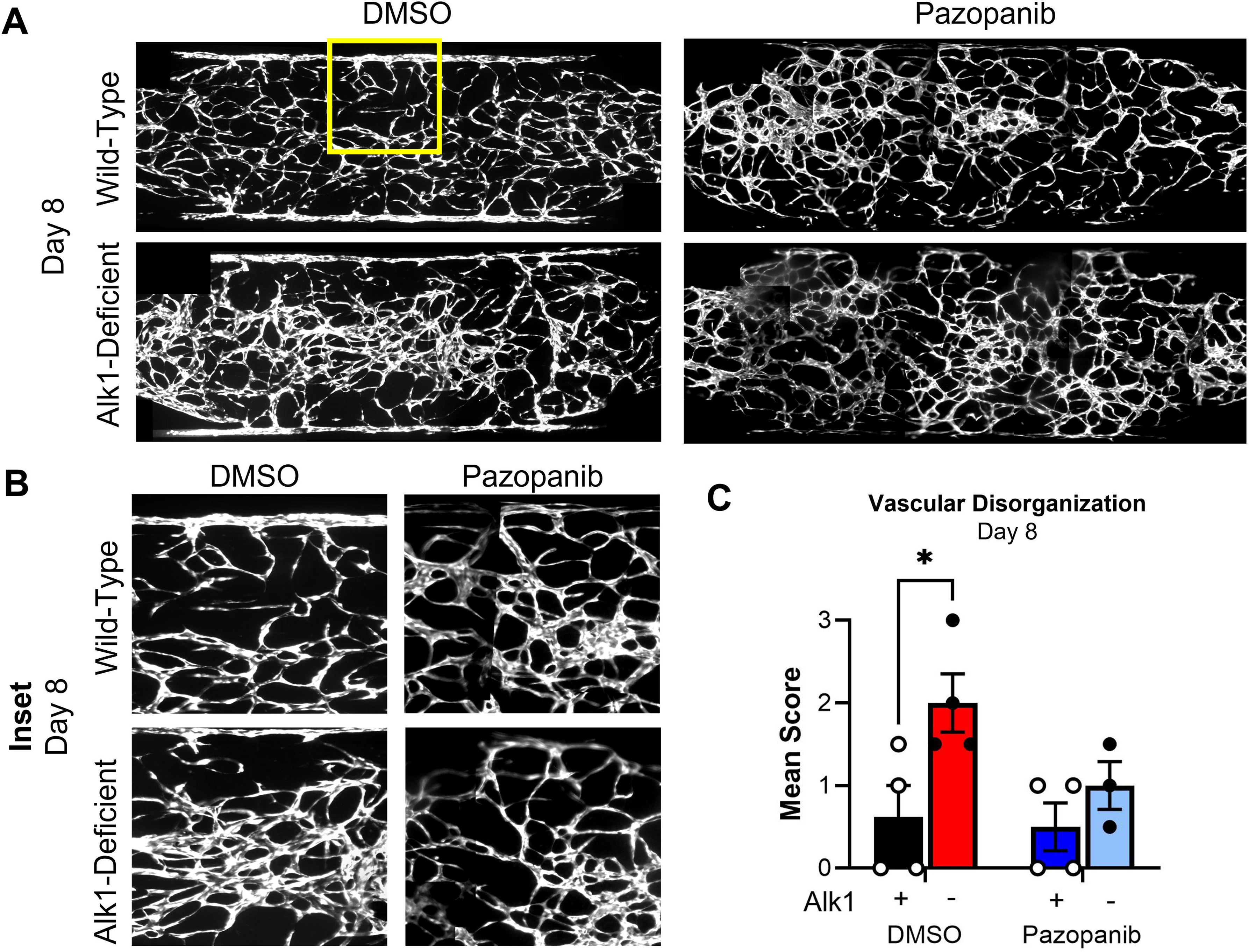
Pazopanib prevents microvascular lesions in HHT-VMO. **A)** IPTG was administered to initiate Alk1 knockdown at day 0, and 100nM Pazopanib (or DMSO) was added to circulating media beginning at day 3. **B)** Pazopanib prevented subsequent appearance of microvascular lesions in IPTG-induced Alk1-knockdown networks. **C)** Vascular disorganization scores are elevated with IPTG-induced Alk1-knockdown in DMSO-treated vehicle control, but not in networks treated with Pazopanib. (Statistics: * - p < 0.05).

### Enrichment for strongly Alk1-deficient EC drives lesions reminiscent of arteriovenous malformations

The approach described above reproducibly produced disorganized and dilated lesions reminiscent of HHT patient telangiectasias, but shunt-like structures did not reliably develop. We hypothesized that this might be because Alk1-deficiency is heterogeneous across the transduced EC population due to the relatively low MOI used (see **Materials and Methods**) along with a normal distribution of lentiviral uptake; thus, there might be incomplete penetrance of Alk1 knockdown. To remedy this situation and to enrich for more consistency in Alk1-knockdown, we used a brief puromycin pre-selection prior to cell seeding in the HHT-VMO (or in a modified version of the platform redesigned to fit a standard 96-well plate format while keeping all other parameters consistent, **Supplemental Figure 8**). To mimic the *in vivo* situation, where only some EC are homozygous mutant due to an LoH event,^12^ we co-seeded wild-type EC along with purified sh-*ACVRL1* EC in a one-to-one ratio. Remarkably, we now observed that on addition of IPTG, we could consistently develop shunt-like structures (**Figure 6**). Conduits were especially obvious when intravascular flow was marked by perfusion of 70kDa fluorescent dextran, which flowed directly from the upper (arteriole-like) to the lower (venule-like) microfluidic channels, through these enlarged vascular conduits (**Figure 6A**).

**Figure 6.**
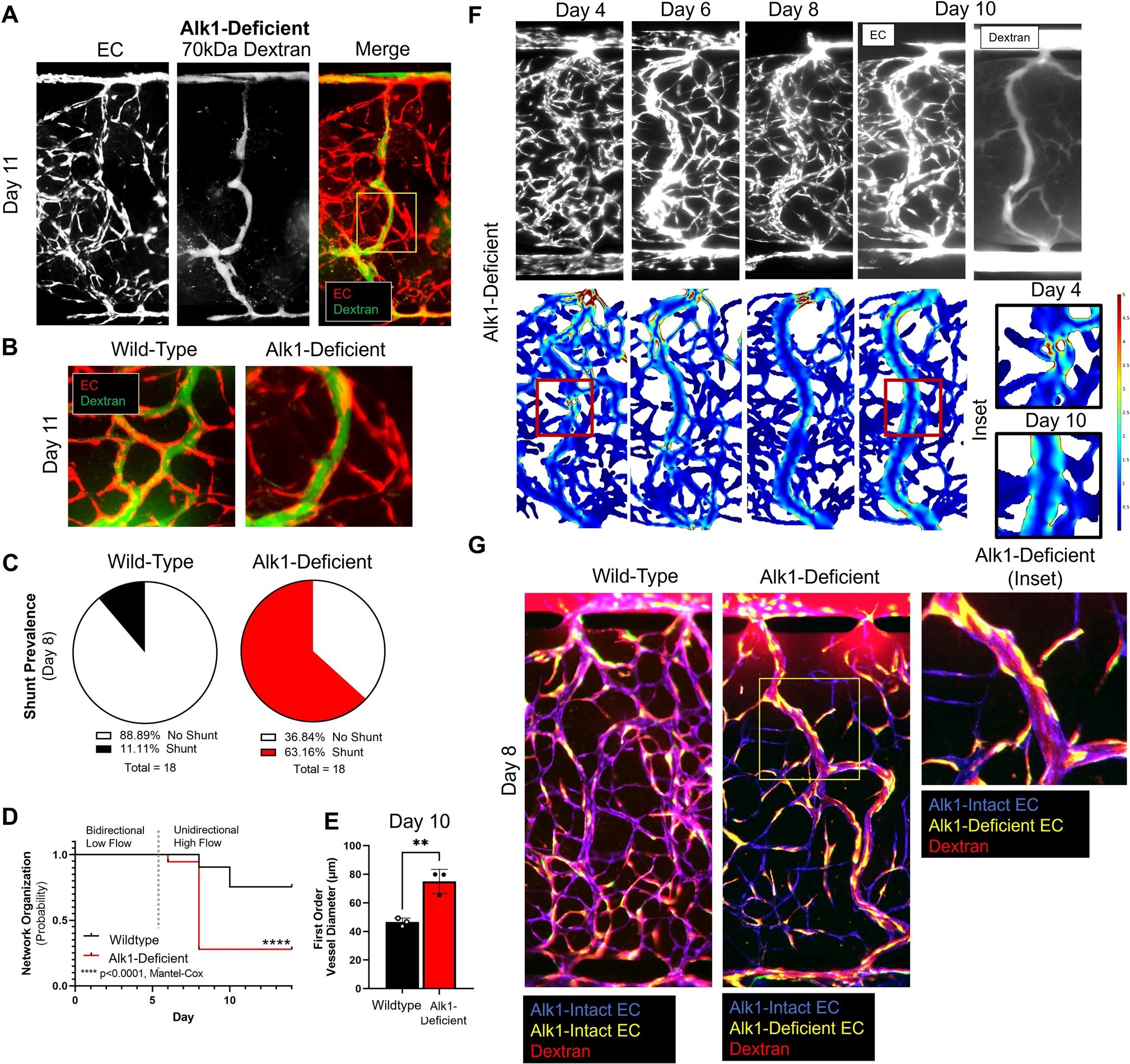
Enrichment for Alk1-deficient EC drives formation of shunt-like structures. **A)** shAlk1 EC were purified using puromycin pre-selection and then co-seeded at a 1:1 mixture with untransduced (Alk1-intact) EC. Under these conditions, Alk1-deficient networks developed enlarged shunt-like structures by day 11 that directly connected upper and lower microfluidic channels. **B)** Intravascular flow (marked by perfusion of 70kDa fluorescent dextran (green)) bypassed adjacent microvessels. **C)** Shunt-like structures were observed in 63.2% of Alk1-deficient networks, compared to only 11.1% of wild-type networks. **D)** Shunt-like structures were evident by day 8, after switching from bidirectional low-flow to unidirectional high-flow conditions. **E)** Mean vessel diameter of shunt-like structures was significantly increased compared to first-order vessels in the same region of wild-type networks (Statistics: ** - p<0.01, Students’ t-test). **F)** *In silico* modeling of intravascular flow revealed that dilated lesions were associated with high shear stress in downstream branching vessels, which subsequently enlarged and appeared to anastomose over time to form the complete shunt. **G)** When Alk1-intact and Alk1-deficient EC were engineered to express different reporter proteins, shunt-like structures were comprised of both endothelial cell types. (Statistics: ** - p < 0.01).

A key feature of a developing AVM is that due to its increasing diameter, it carries a greater fraction of the available blood, and thereby starves adjacent branching microvessels. These branching collateral microvessels eventually regress, leaving an enlarged direct artery-to-vein connection – an AVM – in place of the previous balanced capillary bed. We observed a similar phenomenon in the HHT-VMO, and again, this was particularly apparent when vessels were perfused with fluorescently-labeled dextran. Shunt-like vessels were strongly fluorescent, whereas the adjacent capillary-like vessels became extremely narrow and appeared to cease supporting flow (**Figure 6B**). Disorganized, shunt-like lesions formed in 63.2% of IPTG-treated networks, compared to only 11.1% of control networks (**Figure 6C**), and these vascular structures typically formed by day 8, two days after networks were switched from bidirectional low flow to unidirectional high flow (**Figure 6D**). In the retina of Alk1 mutant mice, shunts develop preferentially in first-order retinal vessels – i.e., vessels that branch immediately from the primary arteries that feed retina – that are exposed to high fluid shear.^36^ Consistent with this finding, we also observed that shunts in the HHT-VMO appeared more reliably in first-order vessels in HHT-VMO networks – i.e., the first branching vessels immediately downstream from the upper microfluidic channel where fluid velocity and shear stress is calculated to be highest (**Figure 2F-G**). These shunt-prone first-order vessels were measured at day 10 for their vessel diameter at the upper, middle, and lower regions of the tissue chamber. In networks containing Alk1-deficient EC, the mean diameter of shunt-like structures was 75.0 ± 19.1 μm, compared to 46.6 ± 11.7 µm in Alk1-intact (i.e., no IPTG) control networks (**Figure 6E**). This corresponds to a 61% increase in first-order vessel diameter in Alk1-deficient vessel networks compared to wild-type controls, and a 20-fold increase in flow, consistent with the development of a shunt.

To better understand the progression by which first order vessels enlarge into shunt-like conduits in Alk1-deficient networks, we imaged fluorescent vessels to track shunt formation over time. At day 4, overall microvascular architecture appears relatively organized, although focal, disorganized telangiectasia-like structures – if not shunts – are already apparent. This is consistent with the early hyperdense phenotype we reported at this timepoint in **Figure 3**. Interestingly, when we used *in silico* modeling to understand how intravascular flow is affected by the presence of these telangiectasia-like lesions, we found that flow (and thus fluid shear stress) was variable throughout the length of these first-order vessels (**Figure 6F, inset**) Specifically, we found that shear is elevated immediately downstream of a dilated, telangiectasia-like tangle (**Figure 6F**, day 4 inset). By day 10, we found that these high-shear regions had outwardly remodeled to increase vessel diameter resulting in reduced and more homogenous shear – a normal vascular response to high focal flow and shear to redistribute flow primarily through these large shunt-like conduits – resulting in an apparent AVM (**Figure 6F, inset**). Further studies are needed to confirm this finding at greater time- and fluorescent-imaging resolution.

### Shunts are comprised of both Alk1-intact and Alk1-deficient EC

A reasonable hypothesis regarding lesion formation in HHT is that a single clone of HHT-mutant EC expands to dominate a region of the vascular network, and that this region undergoes AVM formation. This has been previously observed to drive vascular malformation in diseases such as Cerebral Cavernous Malformation (or, CCM).^40^ To assess this hypothesis, we expressed red or green fluorescent reporters in wild-type (Alk1-intact) and sh-*ACVRL1* (Alk1-deficient) EC populations, respectively, in the HHT-VMO. We applied IPTG (or not) beginning at day 0 to induce Alk1 knockdown selectively in sh-*ACVRL1* EC. For this study, because only rhodamine- or FITC-conjugated dextrans were available, EC were imaged on red and green channels first, and then immediately re-imaged following administration of 70kDa rhodamine dextran to perfusing media to mark intravascular flow. Pre- and post-dextran images were then stitched together, manually overlaid, and differentially false-colored in order to distinguish each (Alk1-intact vs. Alk1-deficient) EC population from themselves and from the perfused fluorescent dextran. In control networks (lacking IPTG) wherein both EC populations (blue and yellow, **Figure 6G**) express Alk1, EC were integrated evenly into the network at day 8 and both telangiectasia-like and AVM-like vascular lesions did not develop. Instead, all microvessels were well perfused as determined by distribution of circulating 70kDa fluorescent dextran (**Figure 6G**). Perhaps surprisingly, and certainly contrary to our initial hypothesis, on addition of IPTG to knock down Alk1 in half of the cells (Alk1-deficient, yellow), we found that AVMs formed and that these were chimeric, compromised of an intermingling of blue (Alk1-intact, wild-type) and yellow (Alk1-knockdown) EC (**Figure 6G and inset**). This has profound implications for the mechanism of shunt formation, as discussed below. Importantly, no aberrant vessel morphology was observed when EC transduced with control vector were co-mixed with wild-type EC (**Supplemental Figure 9**).

### Gene expression changes in Alk1-deficient microvasculature

To better understand gene expression changes in Alk1-deficient EC that may contribute to lesion formation, we generated control networks and mosaic networks comprised of both Alk1-intact and Alk1-deficient EC (as described in **Figure 6**) and performed single-cell RNA sequencing (scRNA-seq) (**Supplemental Figure 10A-D)**. We identified four distinct cell type clusters, including cells with gene expression signatures consistent with EC, pericytes, smooth muscle cells and fibroblasts (**Figure 7A, B**). All four populations were observed in both control and IPTG-treated (Alk1-deficient) networks (**Figure 7A, subset**). Sub-clustering of only EC also revealed four distinct cell clusters (**Figure 7C, Supplemental Fig 10E-I**). Interestingly, cluster 2 was strongly enriched for EC derived from IPTG-treated (Alk1-deficient) networks (**Figure 7D**). Further analysis of cluster 2 EC (i.e., “EC2”) revealed reduced expression of *ACVRL1* (Alk1), as expected, as well as reduced expression of *ENG* (Endoglin) (**Figure 7E**). Significantly, several genes shown to be overexpressed in telangiectasias from HHT2 patients^41^, were also upregulated in EC2, including EMP2, FOSL1, NEXN, AKR1B1, and PHLDA2. We also saw down-regulation of several pro-angiogenic genes, including ANGPT2, KDR, FLT1 and DLL4, which at first seemed surprising given the vascular overgrowth phenotype associated with telangiectasias. However, the scRNA-seq was performed on tissues at the time of AVM formation (see **Figure 6**), where smaller vessels are regressing as flow is diverted to the shunt. Further analysis of the differentially expressed genes (DEGs) in EC2 versus all other ECs (**Supplemental Table 1**) strongly supported the idea that EC2 represents a remodeling population of cells.^42^ For example, we note downregulation of genes associated with vessel stability, such as KLF2, FZD4, NOS3, and TSPAN12^42^ (**Figure 7F**). Concomitantly, we see an upregulation of genes directly associated with vessel regression and pruning, including TAGLN2, CDKN1A, and MAPK10^42–44^ (**Figure 7G**). Several genes associated with matrix and matrix remodeling are also expressed, including FN1, COL1A1, COL1A2, COL3A1, COL5A2, COL6A1, COL6A2, COL6A3, COL8A1, COL12A1, TGFBI, PCOLCE, VCAN, MMP2, TIMP1, TIMP3, SERPINE1, ADAMTS2, ADAMTS12, and ADAM12. In addition, both pro- and anti-angiogenic semaphorins are expressed, notably SEMA3A, SEMA3C and SEMA5A. PDGFB, which is involved in mural cell recruitment to vessels, was also reduced in Alk1-knockdown cells (**Figure 7F**). Consistent with these data, gene ontology (GO) analysis of upregulated genes following Alk1 knockdown (EC2) showed an association with EC migration **(Figure 7H)**. In contrast, GO analysis of upregulated genes in control networks showed an association with establishment of endothelial cell barrier, regulation of angiogenesis, and regulation of vasculature development (**Figure 7H**). Finally, pathway analysis focusing specifically on the PDGF pathway highlighted the loss of PDGFB-PDGFRβ signaling from EC to pericytes and SMC (**Figure 7I**). Thus, EC2 appears to represent a population of cells that are actively remodeling vessels from a stable and complex vascular network to a relatively small number of large AVM-like vessels.

**Figure. 7.**
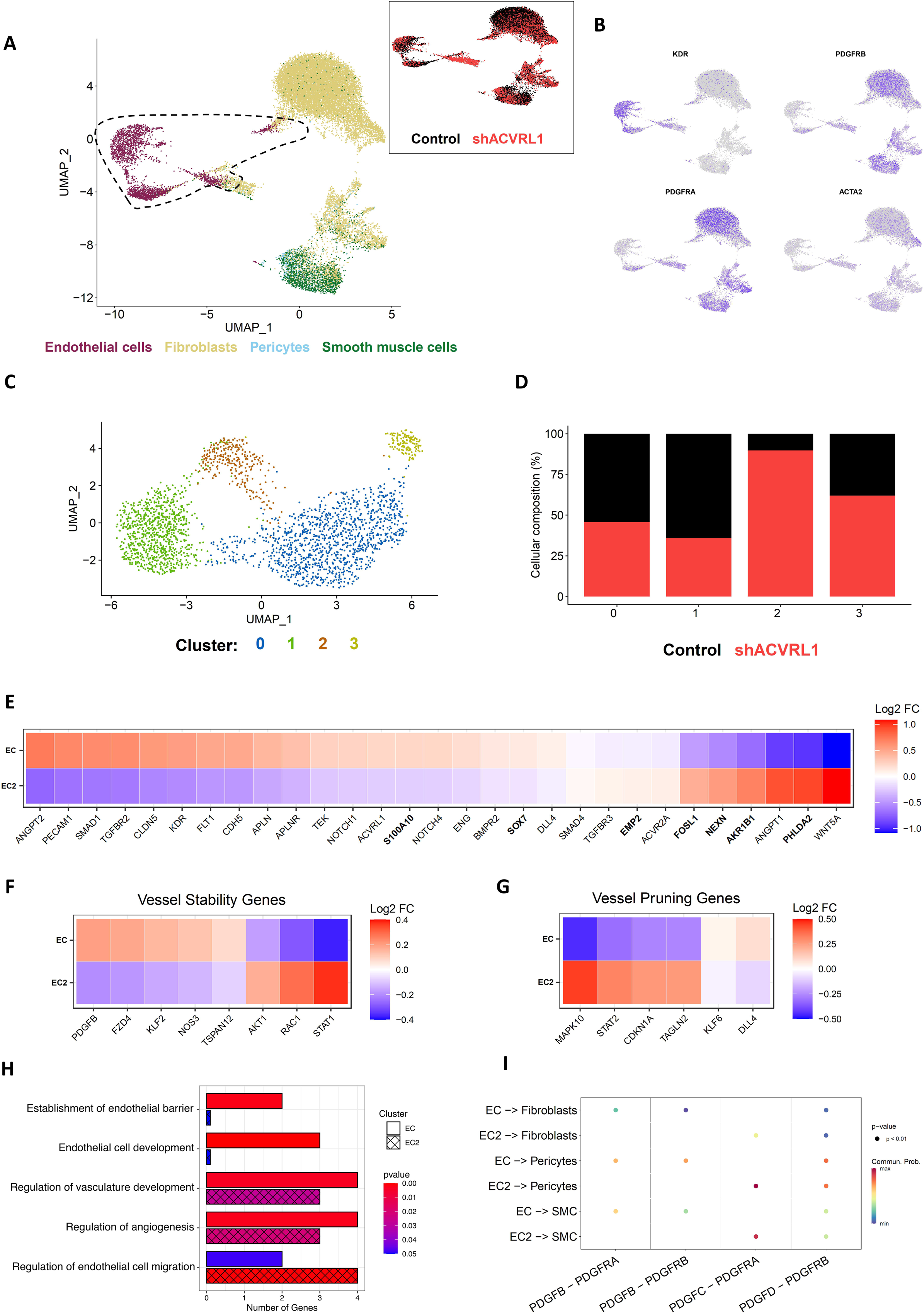
Identifying endothelial cell heterogeneity in control and shACVRL1 vascular micro-organs. Single-cell RNA sequencing of the control and shACVRL1 datasets shows **A)** that each population contains endothelial cells, fibroblasts, pericytes, and smooth muscle cells (subset) with certain regions being unique to either dataset. **B)** Expression of common marker genes distinguishes endothelial cells from the stromal population. **C)** Subsetting of endothelial cells (dashed outline in A) and integration. Unsupervised clustering shows four distinct clusters. **D)** Cluster 2 is mainly comprised of endothelial cells from the shACVRL1 dataset. **E)** Heatmap showing average expression of TGF-β associated genes, angiogenesis-associated genes, and genes from clinical samples (bolded). **F)** Heatmap showing decreased expression of vessel stabilization genes in EC2^42^. **G)** Heatmap showing increased expression of vessel pruning genes in EC2^42^. **H)** Bar plot of pathway analysis on the top DEGs for EC2 and non-EC2, showing the number of genes in the DEGs for each pathway as well as the p-value for the pathway. **I)** Cell-cell communication for EC to stroma evaluating PDGF signaling.

## Discussion

Owing to their low-cost, reproducibility, and ease-of-use, OoaC/MPS technologies offer new avenues for the study of healthy physiology, disease biology, and preclinical drug screening using fully human tissues.^25,45,46^ In particular, vessel-on-a-chip platforms are a potentially powerful tool for studying the pathogenesis of vascular malformation, especially in designs where EC are not constrained to form vessels along pre-established microfluidic channels and structures. For example, in our base vascularized micro-organ (VMO) platform (as well as earlier prototypical iterations of this device), co-seeded human EC and stromal cells freely self-organize in response to interstitial flow to form a perfused microvascular network within a three-dimensional hydrogel microenvironment.^27–30^ Recently, Soon and colleagues^47^ used a similar design to model brain arteriovenous malformations (AVM) by co-mixing in EC expressing mutant variants of *KRAS4*. Resulting networks were abnormal in appearance and included regions of dilated vessels. This study underscores that vessel-on-a-chip technology that incorporates flow can serve as a basis for modeling vascular malformation defects relevant to several disease contexts, including HHT. Indeed, in the present work, we were able to show a hyperdense and abnormal phenotype in the HHT-VMO using both siRNA and shRNA to knockdown Alk1 in primary human EC. However, shunt-like structures did not develop in the base VMO platform, suggesting that even in the presence of Alk1-deficiency, additional signals are required to drive shunt formation.

Recently, Orlova and colleagues seeded HHT1 patient-derived human iPSC-derived (*ENG-*haploinsufficient) EC (iPSC-EC) into a vessel-on-a-chip platform. Resulting networks were distinct in morphology compared to microvessels formed from isogenic control iPSC-EC; surprisingly however, although microvessels were more permeable and fragile, HHT-mutant microvascular networks had reduced (rather than increased) mean vessel diameter and vascular density, suggesting that HHT-like vascular lesions did not develop in this setting. One possible explanation is that the signals that drive HHT-associated vessel malformations (i.e., telangiectasias and shunts) may not be sufficient in some vessel-on-a-chip designs to produce definitive vascular malformations. For example, patient-derived iPSC-EC are heterozygous for HHT-causing gene mutations and retain residual gene expression from their remaining intact allele; as such, a second-hit LoH that ablates residual Alk1 or Eng gene expression – a proposed precursor to lesion formation in HHT patients^11,12^ – may be absent.

We and others^22,36,48^ believe that an additional critical signal for AVM formation in HHT is hemodynamic flow. Using an HHT mouse model, Larrivee and colleagues showed that shunts typically form in high-flow regions of the neonatal mouse retina close to the optic nerve,^36^ where *in silico* modeling predicts wall shear stress values are higher than in the retinal vascular periphery,^37^ and where the retinal vasculature typically undergoes EC quiescence, arteriovenous specification, and vessel organization and maturation.^49,50^ Furthermore, Rochon and colleagues show in zebrafish models that flow-sensitive EC migration is dysregulated with Alk1-deficiency^22^ which may reflect a defect in mechanosensitive signaling transduction in mutant EC. In the current study, we find that gravity-driven flow in our base VMO platform produces shear stress values of ∼0.5 dynes/cm^2^ -- clearly sufficient to support the formation of vessels-on-a-chip,^28^ but apparently insufficient to support the development of AVMs (as evidenced by our finding that Alk1-deficient EC do not generate shunts in the base VMO, **Figure 1** and **Supplemental Figure 1**).

To address these and other issues, we redesigned our existing VMO platform to support a significantly increased gravity-driven intravascular flow, resulting in ten-fold greater wall shear stress values of ∼5 dynes/cm^2^. We also enlarged the tissue chamber to support a broader range of flow and shear stress profiles across the resulting microvascular network. The HHT-VMO design supports microvascular network formation similarly to the original VMO, and higher flow seems to support additional arteriovenous specification. Seeding of Alk1-deficient EC and growth under lower flow conditions produces hyperdense microvessel networks with development of enlarged and dilated focal lesions reminiscent of HHT patient telangiectasias. Further enrichment for more penetrant Alk1-deficiecy in EC drove the development of shunt-like AVM structures under higher flow conditions. Importantly, we demonstrate in several experiments that lesions respond similarly to *in vivo* HHT mouse models. For example, consistent with published work in mice,^19,20^ Alk1-deficiency during periods of active vessel growth produces lesions in the HHT-VMO, whereas loss in established vessels does not. We also showed that treatment with the VEGFR inhibitor pazopanib – which appears to reduce HHT-associated bleeding in patients – prevented the appearance of lesions in Alk1-deficient networks.

An important finding in this study is that lesions are comprised of both wild-type (i.e., Alk1-intact) and Alk1-deficient cells, and that these are seemingly randomly inter-mixed. This is in contrast to a model, with some appeal, whereby a clone of mutant cells expands to dominate the vessel wall, triggering lesion formation, as is reported to occur in Cerebral Cavernous Malformation.^40^ Instead, these data favor a cell non-autonomous mechanism for shunt formation and are consistent with the recent work of Snellings and colleagues^12^ which reported that HHT patient lesions are heterogeneous for both HHT gene-expressing EC and gene-ablated EC (by a proposed second-hit LoH event). These data are also consistent with mouse studies showing that minimal Endoglin deletion in a subpopulation of EC is sufficient to produce a lesion when combined with a pro-angiogenic stimulus.^14^ It is not yet clear how the presence of randomly positioned mutant EC can trigger a vessel-wide lesion, however our gene expression data hint that cell-cell communication may well play a major role as we noted mis-regulation of several junctional and signaling molecules in our single-cell RNA sequencing analysis (**Figure 7)**. Further studies are needed to address these ideas, including by taking advantage of the HHT-VMO’s ability to support real-time observations of lesion formation (as presented in **Figures 3 and 6**). We also see evidence in the scRNA-seq data for a population of EC (EC2) that appear to be associated with vascular remodeling. During shunt formation in the HHT-VMO we see regression of smaller vessels concomitant with the growth of the shunt, which would be consistent with integration of the regressing cells into the larger vessel. Several genes known to be involved in vessel stability or pruning were differentially expressed in EC2 compared to other EC, including KLF2, FZD4, NOS3, TSPAN12,TAGLN2, CDKN1A, and MAPK10. Interestingly, PDFGB, which is a driver of mural cell recruitment to the vessel wall, was also reduced in knockdown cells leading to a loss of PDGFB-PDGFRβ **(Figure 7I)**. Lebrin et al^51^ have previously shown that thalidomide treatment restores mural cell coverage in an animal model of HHT, at least in part through upregulation of PDGFB. Disrupted mural cell (pericyte and SMC) coverage in HHT is thought to relate to the fragile nature of the vessels and their propensity to bleed.^52–54^ However, whether there was a loss of PDGFB expression in the mutant mice was not reported. Finally, the expression of multiple matrix and matrix remodeling genes, along with expression of both pro- and anti-angiogenic semaphorins, marks the EC2 cluster as a unique population of EC, driven by loss of BMP9-Alk1-mediated regulation, that seems to play a critical role in the dissolution of the vascular network and the formation of AVMs.

In summary, the HHT-VMO is an HHT-on-a-chip platform that reliably produces vascular lesions with morphological similarity to the telangiectasias and AVMs that arise in HHT patients. Importantly, our findings match those seen both in mouse models^9,18,20^ and in patient studies.^3,6^ In addition, we have used the model to probe both the timing and the mechanism of lesion formation, and we find that AVM generation is a cell non-autonomous process. Thus, the HHT-VMO captures much of the physiology of intact tissue while offering the increased experimental reproducibility, scale, and ease-of-use benefits of *in vitro* models. Based on this work, we further adapted the HHT-VMO device for a standard microwell plate format, which allows us to integrate readouts that rely on equipment that are fitted for devices with a microplate footprint. Thus, we have significantly increased the scale of studies that can be supported by the HHT-VMO; importantly, mid-throughput drug discovery screens are certainly feasible. We therefore believe that the HHT-VMO can complement and augment existing *in vivo* models of HHT, and that together these tools will further propel research into HHT and the pathogenesis of vascular malformation.

## Materials and Methods

### Cells

HUVEC were selected as our generic EC type for this study because they are easy to isolate and culture, because they retain EC identity for multiplate passages, and because we find that they reliably produce highly organized vascular networks in VMO platforms (e.g., **Figures 1-2**) with a range of vessel calibers within the microvascular range (*data not shown*). We have previously also shown that HUVEC do not endogenously express high levels of arteriovenous specification markers, but are capable of acquiring expression of these specification markers in response to physiological flow.^49^ For this study, primary human umbilical vein endothelial cells (HUVEC) were isolated as previously described,^55^ and expanded in monolayer culture on gelatinized tissue culture flasks in either M199 medium (Thermo #11150067) supplemented with 10% fetal bovine serum (FBS) (Gibco #160000044), 50 µg/mL Endothelial Cell Growth Supplement (ECGS), (Corning #354006) and 50 µg/mL gentamicin (Thermo #15710064); or, in complete EGM-2 (Lonza CC-3162). Primary normal human lung fibroblasts (NHLF, Lonza CC-2512) were cultured in Dulbecco’s Modified Eagle Medium (Corning #10-017-CV) supplemented with 10% FBS (Gicbo) and 50 µg/mL gentamicin (Thermo). All cell lines were routinely tested for mycoplasma contamination, and primary cells were discarded after 8 passages in culture.

### siRNA

HUVEC were transfected with pooled si-*ACVRL1* (Ambion #4392420) or si-*ENG* (Dharmacon #L-011026-00) or non-targeting scrambled control (Dharmacon #D-001810-10 or Ambion #4390843) constructs, as previously described.^49^ In brief, EC were cultured in complete EGM2 over-night prior to transfection (via Lipofectamine 2000, Thermo #11668019) with 8nM siRNA for 24 hours. EC were then washed with fresh complete EGM2 for an additional 24h-48h prior to additional experiments.

### Fluorescent Reporter and shRNA Lentivirus

All lentivirus in this study was packaged with generation II lentiviral packaging vectors in 293T cells, and lentiviral-containing supernatant was tested for mycoplasma contamination and then concentrated in 50% polyethylene glycol (PEG) 8000 (Promega V3011) for at least 72h at 4°C. All lentiviral batches were then tested using HUVEC to calculate concentration of active lentiviral particles. For fluorescent reporter expression, HUVEC were transduced with lentivirus containing CMV promoter-driven red (mCherry, LegoC2), green (eGFP), or blue (Azurite) fluorescent reporters using 8µg/mL polybrene (Fisher # NC9840454) at a multiplicity of infection (MOI) 2 - 5. This typically yields 80-90% transduction efficiency. To achieve inducible Alk1 knockdown, individual *ACVRL1*-targeting short hairpin-RNA (shRNA) clones (**Table 1**) were inserted into the MISSION® 3X-LacO Inducible shRNA plasmid backbone (Sigma). Both clones target a similar region in the *ACVRL1* transcript and produced similar levels of knockdown; thus, both were used interchangeably for this project. sh-*ACVRL1* was transduced into fluorescent EC at an MOI _≈_ 0.8 - 1 in the presence of 8µg/mL polybrene. Using a higher MOI compromised network formation in the VMO or HHT-VMO device, even in the absence of IPTG. Following transduction, EC were recovered from lentiviral transduction with fresh complete EGM2 for a minimum of 24h, at which point they were seeded immediately into the VMO or HHT-VMO; or, for some studies, EC were enriched for high sh-*ACVRL1* transduction by being cultured for 48h in complete EGM2 containing 1µg/mL puromycin (Sigma #8833) and recovered in fresh complete EGM2 for an additional 48h prior to co-seeding into the HHT-VMO device. To knockdown endogenous Alk1 expression, sh-*ACVRL1* expression was induced by addition of up to 3mM IPTG (Sigma #16758) in static monoculture, or in circulating media of VMO or HHT-VMO devices.

### qPCR

Cells in monoculture were washed in PBS and flash-frozen at −80°C or immediately processed for RNA. RNA was isolated in TRIZol™ reagent (Invitrogen) or using a column-based Quick-RNA microprep kit (Zymo R1051) according to manufacturers’ protocols. RNA concentration and purity was quantified using a Nanodrop 2000. cDNA libraries were generated using an iScript cDNA synthesis kit (Bio-Rad #1708890) according to manufacturer protocol. qPCR was performed in a Bio-Rad CFX96 machine using SYBR Green mastermix (VWR #101414) and qPCR primers listed in **Table 2**. All gene expression values were normalized to expression of 18S housekeeping gene.

**Table 2.**
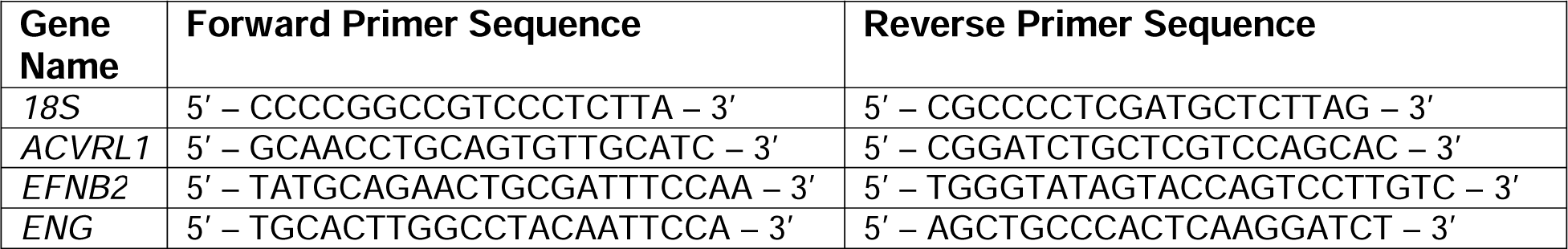
qPCR Primers.

### Immunofluorescence

Adherent cells were cultured in Nunc Lab-Tek chamber slides (Thermo #177399) and fixed with 4% formaldehyde for 15 minutes. Following wash with PBS containing 1% BSA 0.25% Triton X, cells were blocked with PBS containing 3% BSA 0.25% Triton X for 1 hour at room temperature. Cells were then incubated in primary antibody (anti-Alk1 Rabbit Abcam #ab68703, 1:100; anti-CD31 Mouse Dako #M0823, 1:500) over-night at 4°C. Following wash, cells were incubated in secondary antibody (Invitrogen, 1:500) for 2 hours at room temperature. Following wash in PBS, cells were incubated for 15 minutes in 10µg/mL Hoechst 33342, washed in PBS, and imaged on a Leica confocal microscope.

### Western Blot

Cells in monoculture were washed three times in PBS and flash-frozen at −80°C. Cells were scraped into Laemmli buffer containing protease inhibitor cocktail (Roche) and pelleted to remove debris. Protein was quantified using a BCA assay kit (Pierce). 25µg of lysate was run for each sample on a mini-PROTEAN precast gradient gel (Bio-Rad) and transferred onto a PVDF membrane. Membranes were blocked with 5% bovine serum albumin (BSA) TBS-T for 30 minutes at room temperature, followed by exposure to primary antibody (anti-Alk1 Rabbit pAb, Abcam #ab68703, 1:500; or, anti-Tubulin Rabbit Ab, Cell Signaling #2144S, 1:500) overnight in 1% BSA TBS-T at 4°C. Primary antibody was washed in TBS-T and incubated in secondary antibody (Goat anti-Rabbit HRP, Thermo #31460, 1:1000) for 2 hours at room temperature. Following additional wash in TBS-T and TBS, membranes were developed using SuperSignal™ West Femto Maximum Sensitivity Substrate (Thermo #34094) and immediately imaged using a Biorad Geldoc imager. Alk1 signal was assessed first, and then the membrane was stripped (Restore™ Stripping Buffer, Thermo #21059) and probed for expression of Tubulin housekeeping gene expression.

### VMO and HHT-VMO Devices

The vascularized micro-organ (VMO) microfluidic platform^27,28^ was fabricated and loaded as previously described, or adapted for improved study of vascular malformation in HHT (HHT-VMO). In brief, the VMO and HHT-VMO device features were etched onto a silicon wafer using ultraviolet-based photolithography, and then soft lithography techniques were used to generate a microfluidic feature layer in polydimethylsiloxane (PDMS) (Ellsworth, Dow Sylgard 184 #4019862). The features were then enclosed by plasma bonding (Harrick) to a thin commercial PDMS membrane (Pax). For VMO, the enclosed feature layer was plasma bonded (Harrick) to a (3-mercaptopropyl)trimethoxysilane-treated bottomless 96-well culture plate. For HHT-VMO, the feature layer was either PDMS stamped onto bottomless 2mL cryovials (VWR #75852-324) or a custom 3D-printed reservoir layer produced out of high-temperature resin using a Formlabs 3D printer (**Supplemental Figure 8**). Devices were then sterilized using either an autoclave or exposed to 20 minutes of ultraviolet light. Primary EC (less than passage 8) and NHLF (less than passage 10) were resuspended at cell densities of 8×10^6^ cells/mL for each cell type into a hydrogel comprised of 6.5mg/mL fibrin (Sigma #341573) and supplemented with 0.2mg/mL fibronectin (Sigma #F1141) and 10µg/mL aprotinin (Sigma #A-6012). 1mg/mL thrombin (Sigma T4648) was mixed into the hydrogel, and the hydrogel mixture was then immediately loaded into the VMO or HHT-VMO tissue chambers. Upper and lower microfluidic channels were then coated with extracellular matrix protein solution (0.5mg/mL fibronectin, 0.5mg/mL laminin (Fisher 23017015)) for 10 minutes before washout and establishment of gravity-driven complete EGM-2 media circulation through the device. Low interstitial flow (∼2 dynes/cm^2^) of complete EGM-2 was applied such that it alternated direction across the tissue chamber every other day for 6 days, after which point unidirectional high flow (∼10 dynes/cm^2^) was maintained for the remainder of the experiment. Vessel architecture was recorded every two days using an inverted epifluorescent microscope (Olympus or Zeiss). Vessel patency and intravascular flow was confirmed between days 6-8 by addition of sterile 50µg/mL 70kDa FITC- or rhodamine-conjugated dextran (Sigma #FD70S or #R9379) to the media reservoirs, and imaged in microvessel networks of the tissue chamber for 5 - 15 minutes. Fluorescently-conjugated dextran was then washed out by addition of fresh complete EGM2 to the media reservoirs.

### Vessel Morphometry

VMO and HHT-VMO tissue chambers were imaged at 4x or 10x magnification every two days following initial device loading. For HHT-VMO, multiple images were taken across the tissue chamber, and either automatically or manually stitched using Fiji/ImageJ or Adobe Photoshop to capture the entire network. For morphometric analysis, networks were then cropped to a standard size (VMO: 800 x 1200 pixels; HHT-VMO: 1200 x 4000 pixels) to remove microfluidic channels and segmented using the Trainable Weka Segmentation plugin (FIJI/ImageJ).^56^ Segmented images were then analysed using the AngioTool plugin (FIJI/ImageJ)^57^ to obtain vessel area, vessel length, branchpoint number, and mean lacunarity values. For mean vessel diameter across the entire network, FIJI/ImageJ was used to automatically overlay a 10 pixel x 10 pixel grid overtop each segmented network image, identify those grid intersects that coincide with a microvessel, and to measure the corresponding vessel diameter at that point; any grid intersect that fell upon a vessel segment that had already been measured at an adjacent point was excluded from analysis so as not to oversample enlarged vessel structures. For mean vessel diameter across first order vessels, FIJI/ImageJ was used to divide first order vessel structures int top, middle, and bottom segments and to manually measure vessel diameter at evenly spaced intervals within each section. Those measurements were then averaged to obtain vessel diameter across the entire vessel length. For vessel disorganization score analysis, segmented image networks were de-identified using a randomly-assigned five-digit image code. Three investigators were trained using a training image set to assign codes based on morphological features of network architecture, as described in **Supplemental Figure 11**. In brief, higher scores were associated with abnormal network appearance and development of vascular lesions. De-identified experimental image sets were then presented to at least two blinded investigators. In cases where scores from both investigators were highly variable from one another, investigators were re-presented with a de-identified image set that also included some training images interspersed within the dataset. If an investigator failed to reproduce their own scores from training, their scores were excluded from further analysis. Otherwise, investigator scores were averaged for each network once images were reidentified.

### In Silico Flow Modeling

Image masks were generated as described above, and then were binarized, skeletonized, and traced using a custom MATLAB script. The traced images were converted to .DXF files using the DXFLib MATLAB package^58^ and imported into AutoCAD to overlay the schematics for the microfluidic devices. The traced vessels and microfluidic devices were imported into COMSOL 5.2.1, and the velocity and shear stress were calculated using the laminar flow steady-state model. Using the Bernoulli equation, all pressure heads were calculated based on medium height in the inlet/outlet wells and used as input to model the gravity driven flow in the device.

### Pazopanib

Pazopanib (MCE #HY-10208) was resuspended in DMSO at a 10mg/mL stock concentration and added to the HHT-VMO cell culture medium at 100nM concentration beginning on day 3. Cell culture medium containing pazopanib (or DMSO only) was refreshed every two days.

### Single-Cell RNA Sequencing Sample Preparation

The thin transparent membrane was carefully removed from the bottom of the PDMS layer to expose the tissue chambers 12 device units. The chambers were washed with HBSS prior to adding digestion buffer (complete EGM2 supplemented with 400U/mL nattokinase and 1mg/mL collagenase type I) in a drop-wise manner. After incubation at 37°C for 5 minutes and gentle mechanical dissociation with a micropipette, cells suspensions were collected and centrifuged (340 x *g* for 3 min). Cell pellets were resuspended in complete EGM2 at 1000 cells/mL and viability was checked using Trypan Blue to confirm >95% viability. The cellular suspensions were then loaded onto a Chromium Single Cell Instrument (10X genomics) and processed to generate cDNA using 10X Genomics v2 chemistry according to the Chromium Single Cell 3’ Reagents kits v2 user guide.^32^ After undergoing quantification and quality control of cDNA libraries using the Qubit dsDNA HS Assay kit (Life Technologies Q32851), high-sensitivity DNA chips (Agilent 5067-4626), and KAPA qPCR (Kapa Biosystems KK4824), cDNA were sequenced on an Illumina HiSeq4000 at UCI’s Genomics Research and Technology Hub to achieve an average of 50,000 reads per cell.

### Single Cell RNA Sequencing Alignment and Processing

FASTQ files were aligned to GRCh38 and converted to a count matrix with the 10x Genomics CellRanger v6.0.0^59^ using the 10x Genomics Cloud Analysis. The filtered count matrices were loaded into RStudio (R version 4.0.4)^60^ using the Seurat::Read10X_h5 function and converted to a Seurat object using the Seurat R package (Seurat_4.0.0)^61^ with the Seurat::CreateSeuratObject function. Cells were removed if they contained greater than 15% mitochondrial RNA or had fewer than 200 features. In addition, a subset of ribosomal and mitochondrial RNA was removed to improve clustering. Data were normalized and log-transformed using the Seurat::NormalizeData and Seurat::FindVariableFeatures(nfeatures=2000, selection.method = “vst”) functions. The influence of the cell cycle between cycling and non-cycling cells was minimized by regressing the difference between the expression of cells in the S phase and G2M phase using the Seurat::ScaleData function. The data was also scaled to create data in the SCT data slot of a Seurat object using the Seurat::SCTransform function. Doublets were removed using DoubletFinder_2.0.3.^62^ The optimal number of dimensions was determined using the Seurat::ElbowPlot function. The data was simplified to two dimensions using the Seurat::RunUMAP function, and the nearest neighbors were found with the Seurat::FindNeighbors function. The resolution that drives cluster grouping was determined by using the clustree::clustree function^63^ to assess the stability of the clusters, as well as looking at the uniqueness of the differentially expressed genes. The cell types were determined using the SingleR^64^ package and limiting the database to only include cells closely related to those in the devices (endothelial cells, fibroblasts, pericytes, and smooth muscle cells). The control dataset contained 11,224 cells with 1,289 endothelial cells (ECs), 8,739 fibroblasts, 136 pericytes, and 1060 smooth muscle cells (SMCs). The sh-*ACVRL1* dataset contained 8,541 cells with 1,197 ECs, 6,185 fibroblasts, 195 pericytes, and 964 SMCs.

### Single Cell RNA Sequencing Dataset Integration and Subsetting

Datasets were integrated using the scMC^65^ R package, which creates a shared reduced dimensional embedding of cells corrected for technical variation, with the input being the “RNA” slot from the already processed individual datasets. Numerous dimensions were evaluated to test the stability of each. The labeled cell types determined the clustering. As flagged by SingleR,^64^ the ECs for each dataset were subset and then integrated using scMC. The standard Seurat workflow was run to generate unsupervised clusters. Finally, the proportions of the datasets in each cluster were determined using the scMC::computeProportion function.

### Single Cell RNA Sequencing, Differentially Expressed Genes, Pathway Analysis, and Cell-Cell Communication

Differentially expressed genes (DEGs) for cluster 2 of the EC population compared to all of the other EC populations were determined using the Seurat::FindMarkers() function with the DESeq2 technique. The DEGs were filtered by log fold change and percent expression. The top 30 genes were used for pathway analysis. The pathway analysis was conducted using the EnrichR^66^ package with the GO Biological Processes 2021 dataset.^67,68^ Significant pathways related to endothelial cell function were highlighted in the bar plot.

### Statistics

Power analysis indicated that VMO or HHT-VMO experiments require a minimum of 3 replicates are needed to detect significant differences by morphological analysis; thus, a minimum of 3 replicates were included for all studies. Significant differences were detected using Behrens-Fisher t-test, or one-way ANOVA followed by post-hoc t-test for studies involving multiple comparisons. For non-parametric comparisons, a Mantel-Cox rank-sum test was used. In all cases, alpha was set at 0.05.

### Data Access

Analyzed data from this study will be made available through the Biosystics database (https://biosystics-ap.com/), which was formerly known as the Microphysiological Systems Database created by the University of Pennsylvania;^69^ the Gene Expression Omnibus (https://www.ncbi.nlm.nih.gov/geo/) (Accession #: GSE252666); and through figshare (https://figshare.com/), a general data repository. All other data may be made available upon request to the corresponding author.

## Author Contributions

Most of the data were collected and analyzed by JSF, with additional contributions by JA, WVT, DJJ and SM. Single-cell sequencing analysis was performed by CJH. Adaptation of the HHT-VMO for microplate format was conducted by WVT in collaboration with YHC and APL, who were also responsible for portions of device fabrication workflow. CCWH and APL provided project oversight and scientific input. The manuscript was written by JSF, CJH, and CCWH.

## Acknowledgements and Funding Sources

The authors would like to acknowledge all members of the Hughes lab and the Lee lab for their scientific input and technical support. The primary author would like to acknowledge JEL for his help in optimizing the vascular disorganization scoring schema, and for general support throughout the timeline of this study. This project was funded by the following grants: NIH UG3/UH3 TR002137 (Hughes), NIH T32HL116270 (Hughes, Hatch), DOD PRMRP W81XWH2110108 (Fang), a Tulane Committee on Research (COR) Fellowship (Fang), and funds from the Tulane School of Science & Engineering (Fang).

## Notes

### Competing Interest Statement

Author CCWH is co-founder and CSO of Aracari Biosciences, a vessel-on-a-chip biotech company. Aracari Biosciences did not sponsor or fund this project.

### Summary of Updates

Figure 7 was corrected as panels H and I were accidentally from the original version upload.

