## Supplemental Figures for "A Microphysiological HHT-on-a-Chip Platform Recapitulates Patient Vascular Lesions"

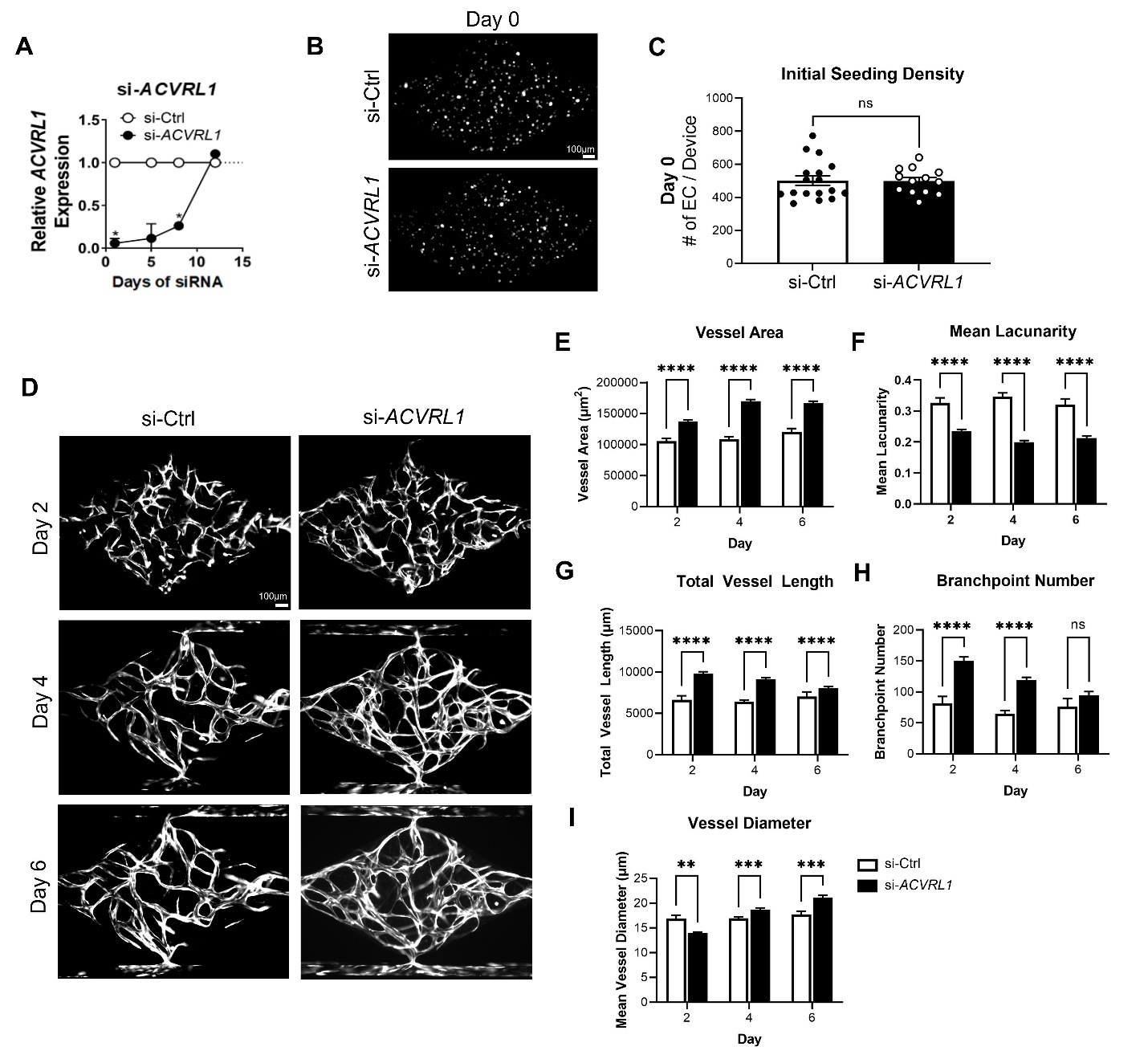

**Supplemental Figure 1. Alk1-deficient EC (via siRNA) form hyperdense microvasculature in VMO.** **A)** Pooled si-*ACVRL1* eliminates >80% endogenous Alk1 mRNA expression in human EC for up to 7 days (vs. si-Ctrl) but knockdown recovers by day 12. **B-C)** siRNA-treated EC are seeded into the VMO platform at similar cell densities. **D)** si-*ACVRL1* form hyperdense microvasculature (vs. si-Ctrl), with **E)** increased vessel area and **F)** decreased mean lacunarity, as well as increased **G)** total vessel length, **H)** branchpoint number, and **I)** mean vessel diameter at most assessed timepoints. (n=16 si-Ctrl, n=13 si-*ACVRL1*)

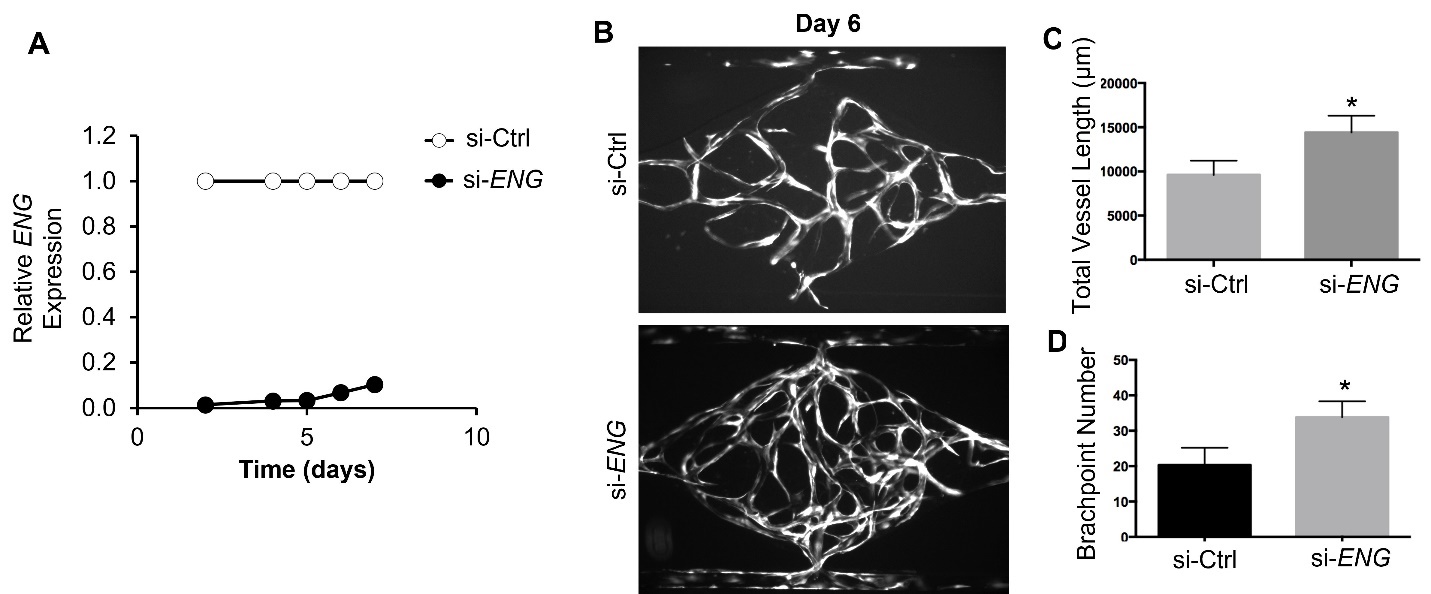

**Supplemental Figure 2. siRNA Knockdown of Eng produces hyperdense microvasculature in VMO.** **A)** Pooled si-*ENG* eliminates >90% endogenous Alk1 mRNA expression in human EC for up to 7 days (vs. si-Ctrl). **B-C)** siRNA-treated EC are seeded into the VMO platform at similar cell densities. **D)** si-*ENG* form hyperdense microvasculature (vs. si-Ctrl) at day 6, with increased **E)** total vessel length and **F)** branchpoint number.

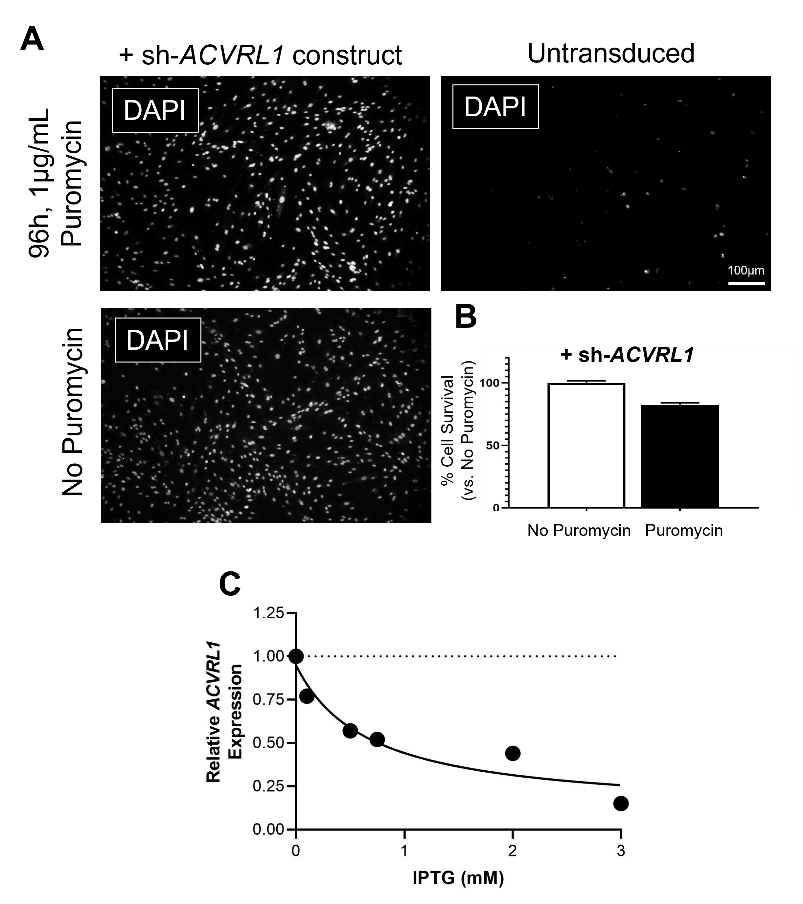

**Supplemental Figure 3. IPTG-inducible sh-*ACVRL1* targets endogenous Alk1 expression.** **A)** Transduction of EC with sh-*ACVRL1* construct confers resistance to 1µg/mL puromycin (96h), whereas widespread cell death is observed in untransduced cells. **B)** ~85% of transduced EC survive puromycin selection after 96h. **C)** Alk1 mRNA knockdown is dependent on IPTG concentration.

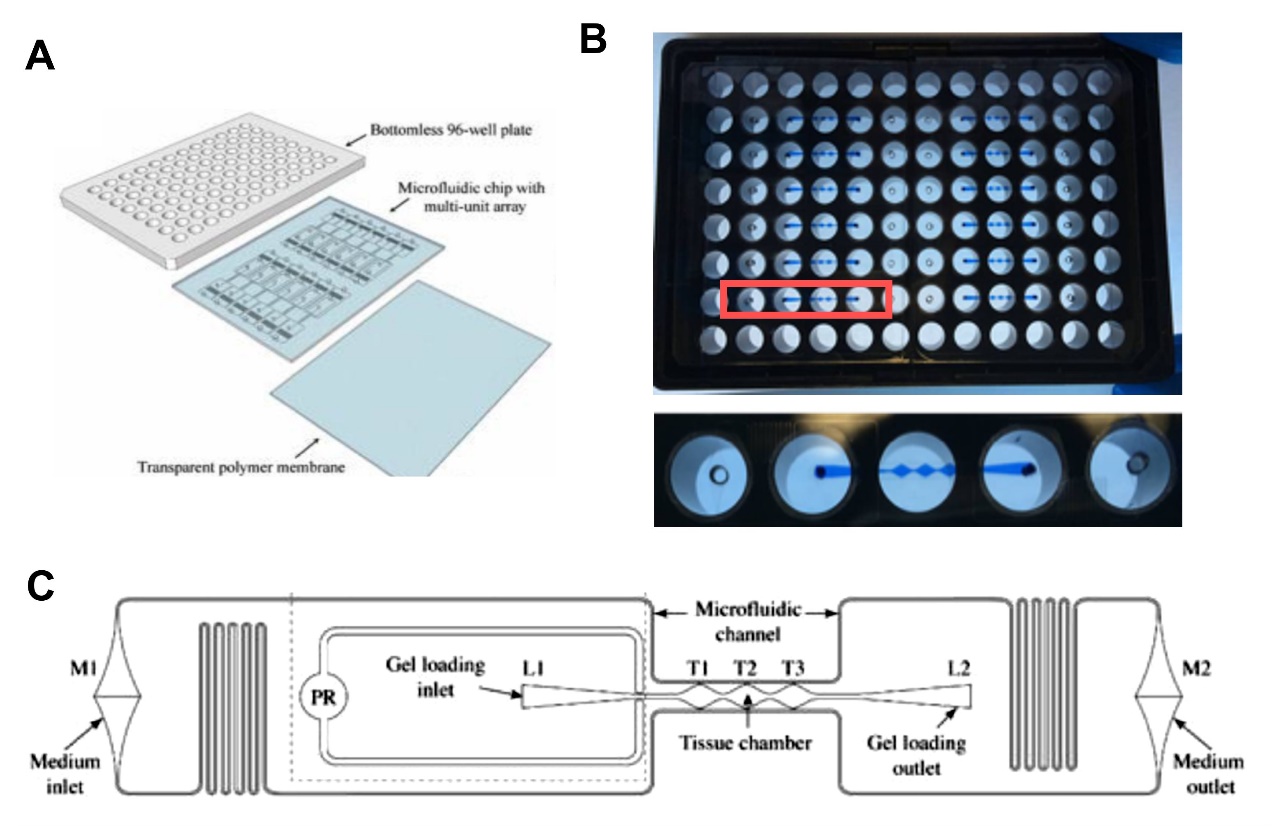

**Supplementary Figure 4: The base Vascularized Micro-Organ (VMO) platform.** As previously published, **A)** the VMO consists of a microfluidic feature layer sandwiched between a bottomless 96-well plate and a transparent polymer membrane. **B)** Each plate contains 8-12 individual bioreactors, each composed of **C)** three central diamond-shaped tissue chambers (T1-3) connected by coupled upper and lower microfluidic channels that circulate media from the medium inlet (M1) and outlet (M2) by gravity-driven flow. (Panels A and C are adapted from Sobrino et al. 2016 and Phan et al. 2017).

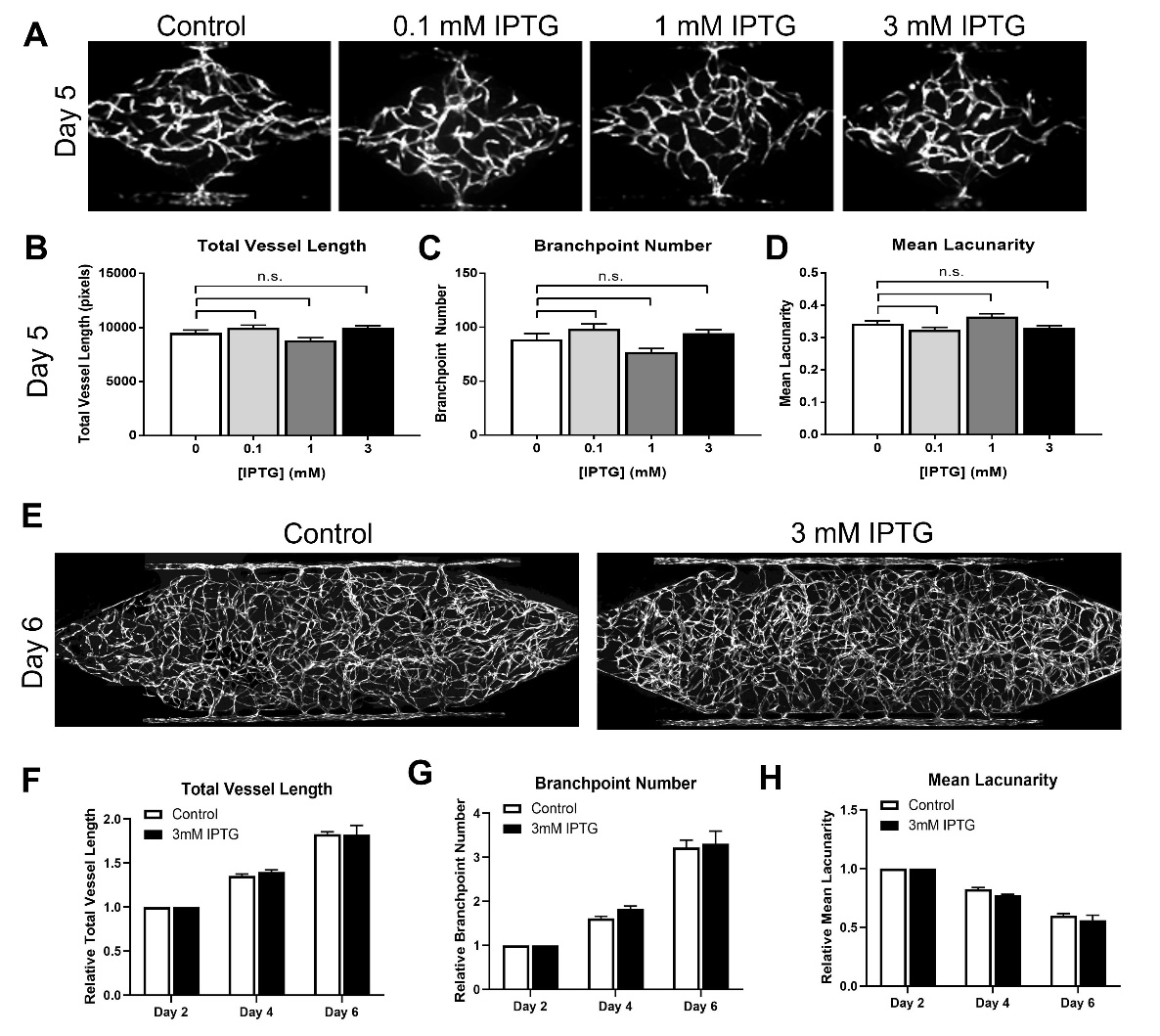

**Supplemental Figure 5. IPTG does not impact microvessel formation or appearance in untransduced EC.** **A)** Addition of 0.1mM – 3mM IPTG to circulating media in the VMO beginning at day 0 has no effect on formation of a microvascular network from untransduced EC. No significant differences were observed in **B)** total vessel length, **C)** branchpoint number, or **D)** mean lacunarity. **E)** 3mM IPTG also did not alter microvessel network formation from untransduced EC in the HHT-VMO platform. No significant differences were observed in **F)** total vessel length, **G)** branchpoint number, or **H)** mean lacunarity.

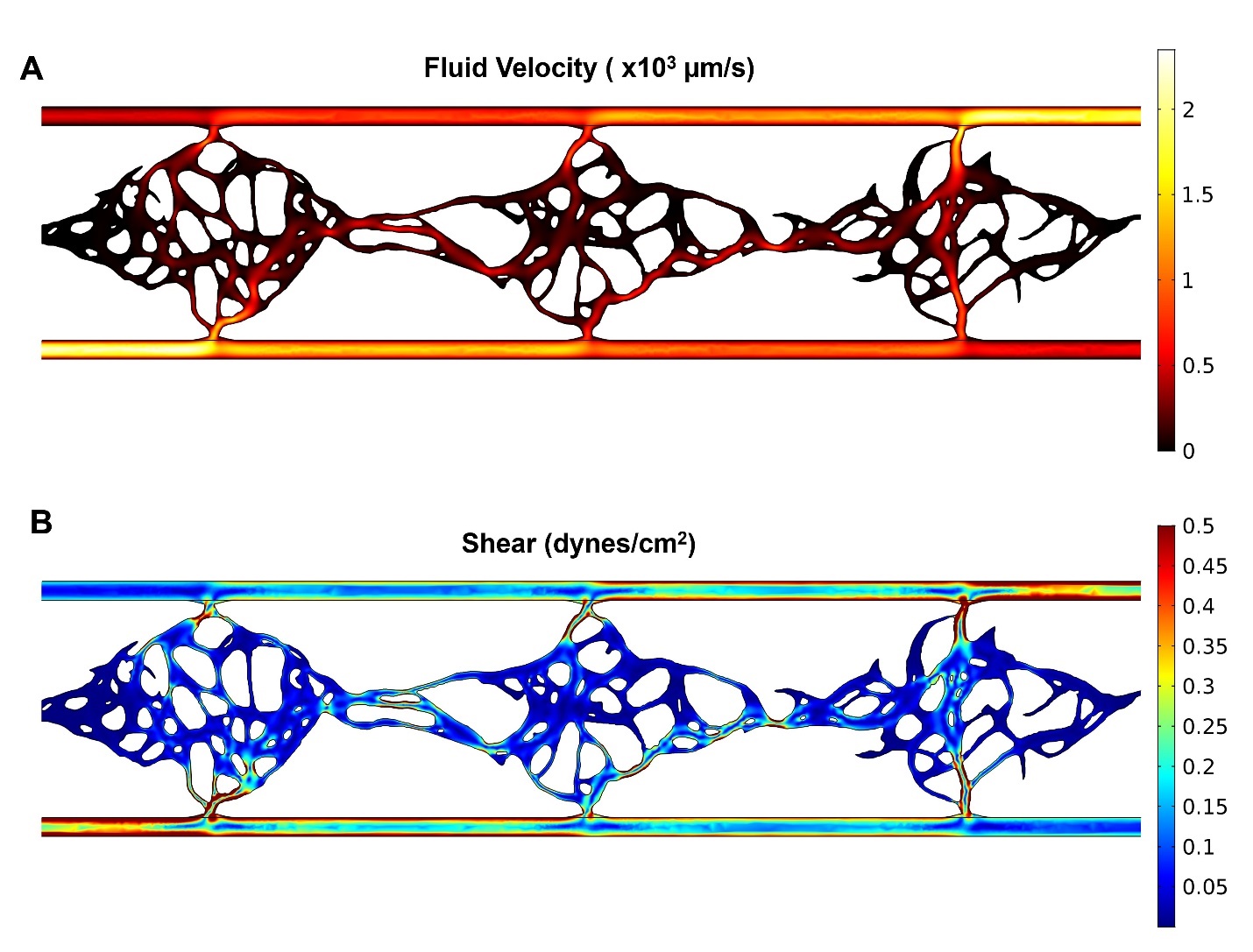

**Supplemental Figure 6. *In silico* modeling of intravascular fluid flow in the VMO.** Established microvascular networks formed from untransduced (wild-type) EC in the VMO were modeled for **A)** gravity-driven fluid velocity (μm/s)and **B)** resulting fluid shear stress (dynes/cm^2^).

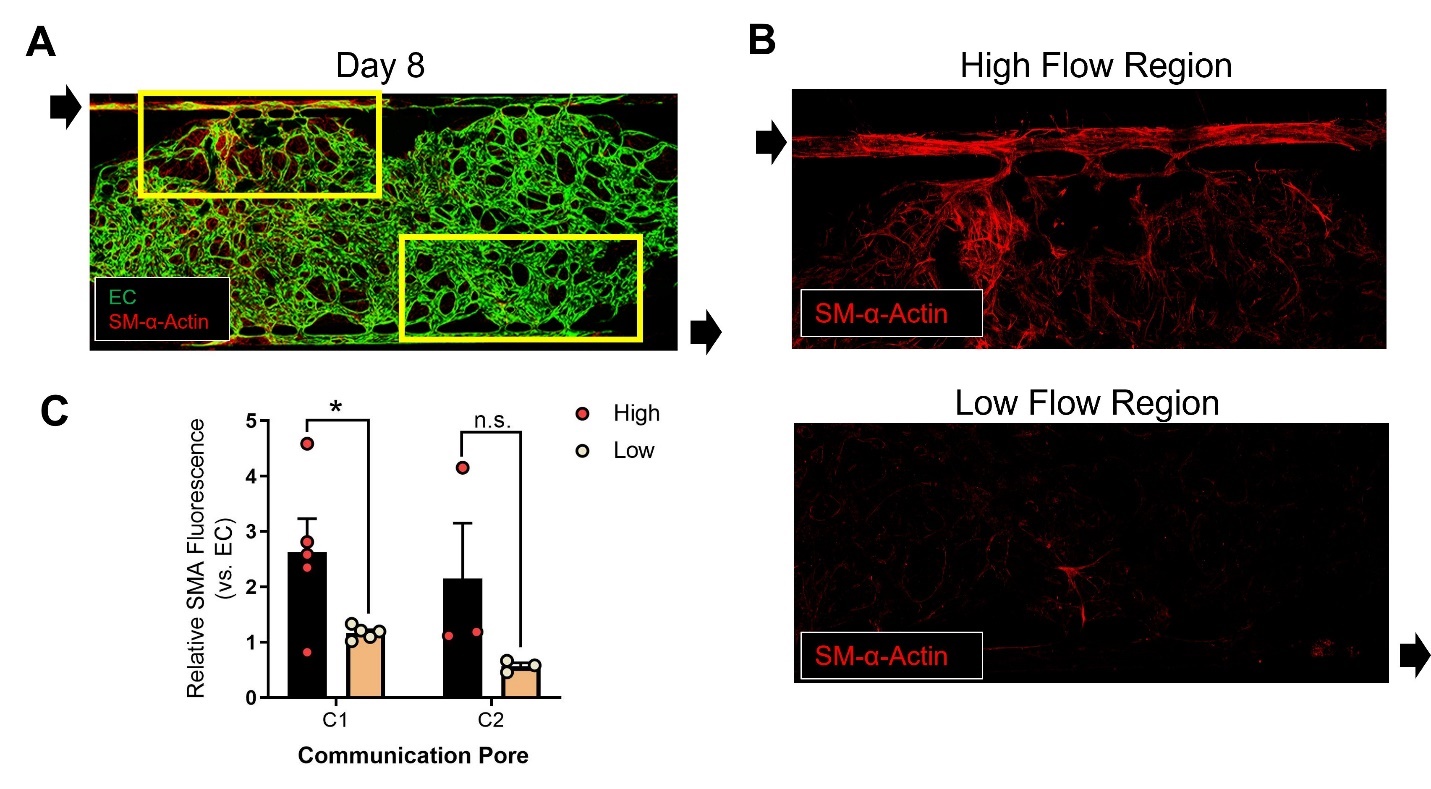

**Supplemental Figure 7. Physiological high flow induces perivascular cells to express SM-α-Actin**. **A)** Exposure of an established microvascular network to physiological high flow for 48h **B)** induces expression of SMA (red) in high flow (but not low flow) regions. **C)** Relative SMA fluorescence (normalized to EC signal) was significantly increased at the first communication pore (C1) of the high flow region (relative to C1 in the low flow region).

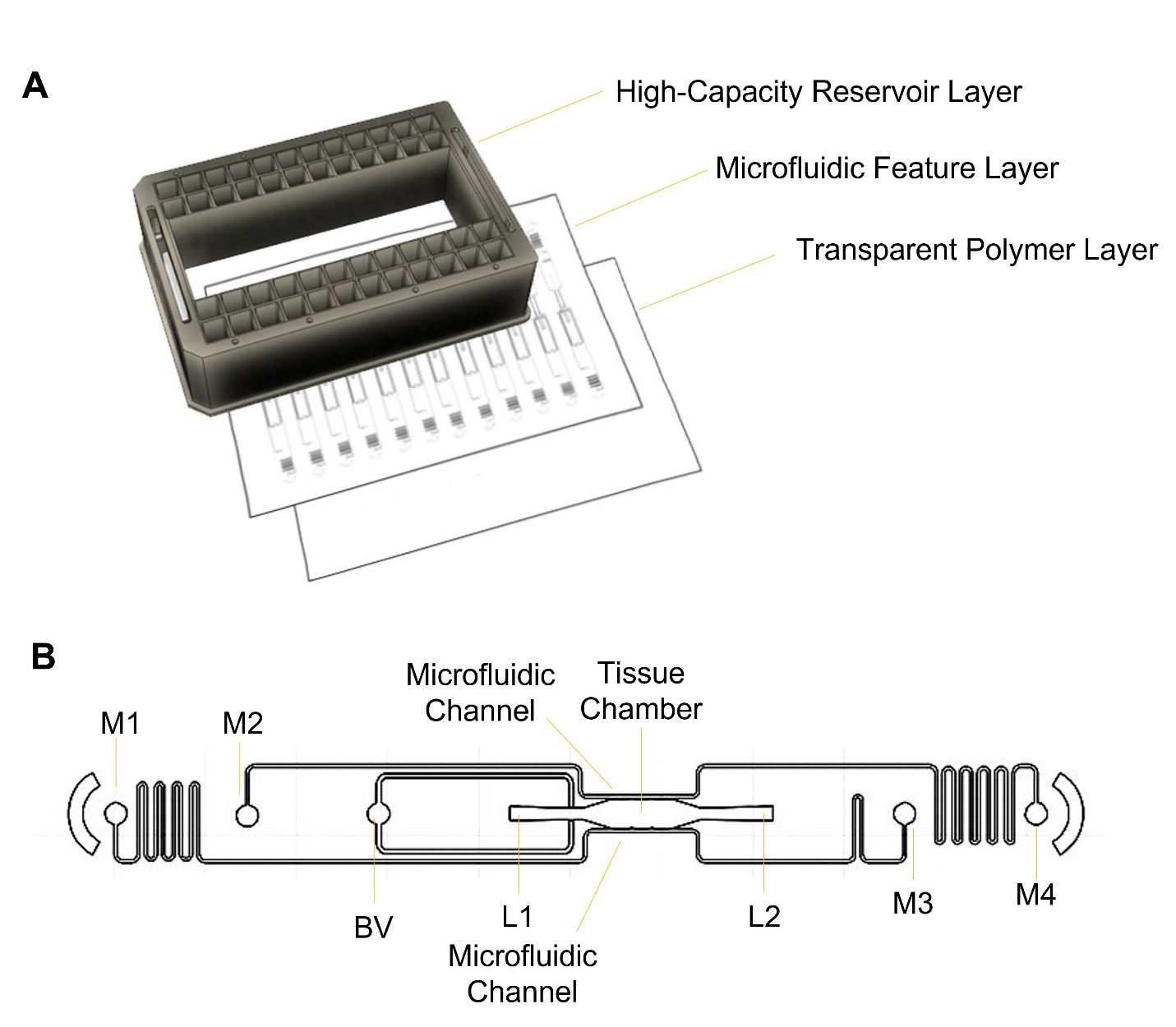

**Supplemental Figure 8: Microplate-adapted versions of the HHT-VMO microfluidic platform. A**) Microplate-adapted HHT-VMO was configured to fit a standard 96-well plate format. It is comprised of a 3D-printed high-capacity reservoir layer that fits a standard microplate lid, a middle microfluidic feature layer, and a transparent polymer layer. **B**) Each feature layer fits 12 microfluidic bioreactors that consist of a central tissue chamber lined by upper and lower microfluidic channels. The chamber is loaded through loading ports (L1, L2), and aided by the presence of a burst valve (BV). Microfluidic channels are connected to media inlets and outlets (M1 – M4) which circulate media contained in high-capacity reservoir wells.

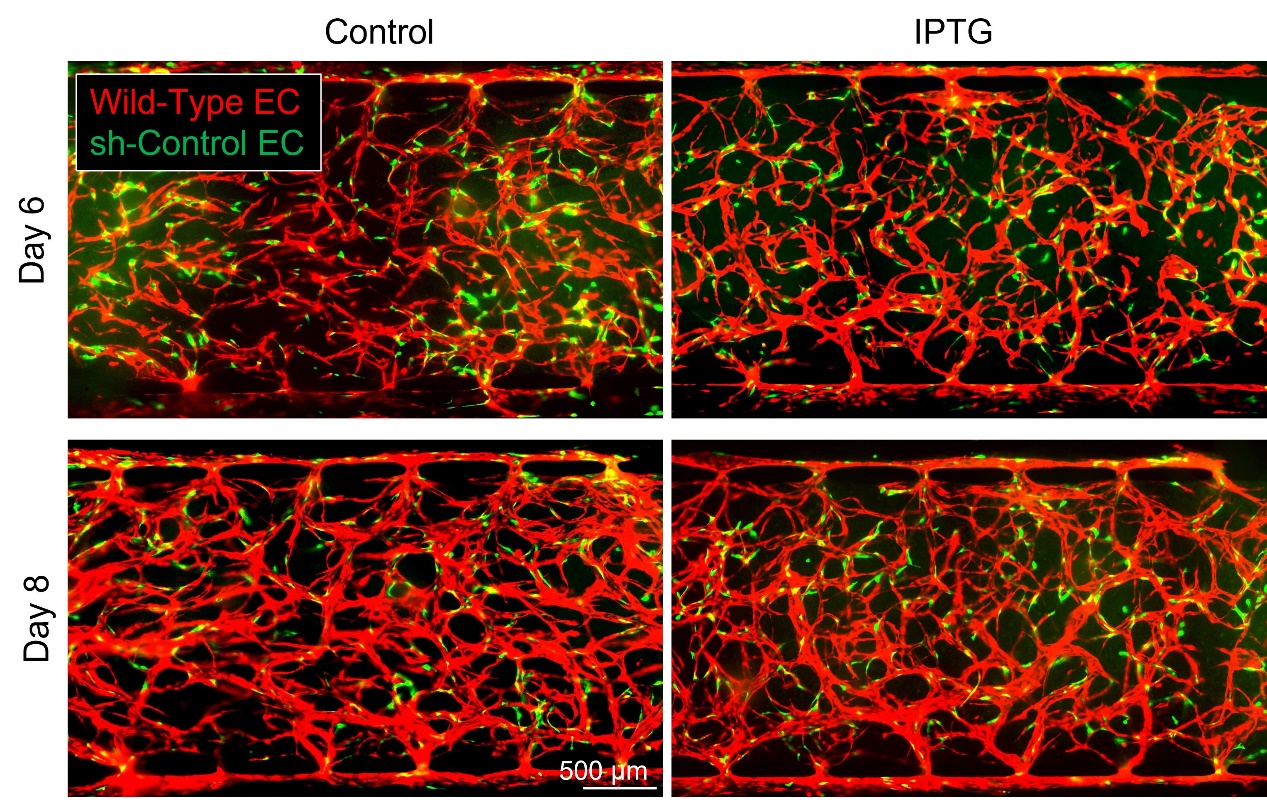

**Supplemental Figure 9: Presence of sh-Ctrl EC do not induce lesion formation in HHT-VMO.** EC transduced with IPTG-sensitive control vector (green) were co-mixed with wild-type (untransduced) EC (red) and seeded into the HHT-VMO. Network appearance remained ordered and normal in appearance at day 6 or day 8, regardless of presence of IPTG in the circulating media, and no shunt-like lesions appeared to develop. (Images representative of n=3 for each group.)

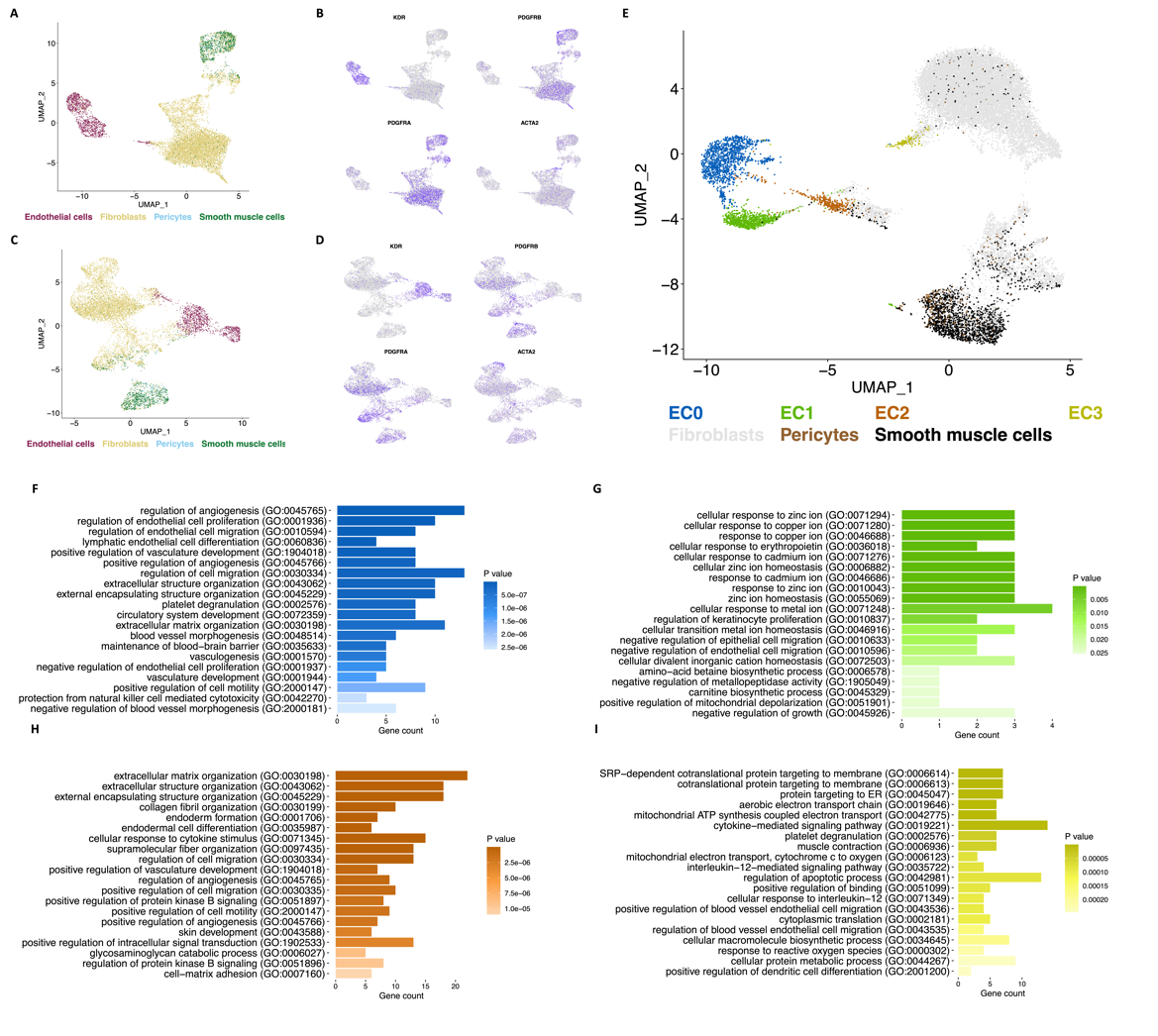

**Supplemental Figure 10. Single-cell RNA sequencing. A-D)** Single-cell RNA sequencing of the **A)** control dataset and **C)** shACVRL1 dataset showing that each are comprised of **B,D)** endothelial cells, fibroblasts pericytes, and smooth muscles with expression of marker genes shown. **E)** Integrated datasets with labeling for cell-cell communication. EC are labeled as cluster 0-3 from assignment based in Fig 7C. **F-I)** Pathway analysis for each EC cluster highlighting different transcriptional changes.

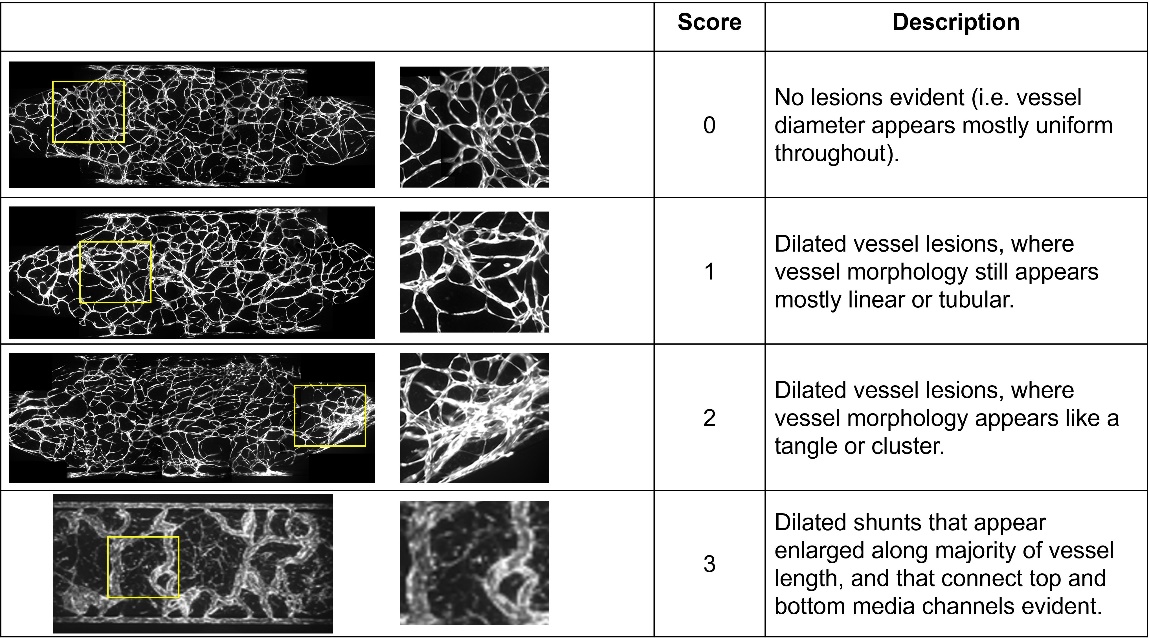

**Supplemental Figure 11: Vessel Disorganization Score.** Scoring table and training images used for vessel disorganization score analysis. (The training image used to represent a severe phenotype – shunt-like lesions - was taken by a different investigator at low magnification and thus left- and right- margins of the tissue chamber were not acquired for this network.)
